# Secondary reversion to sexual monomorphism associated with tissue-specific loss of *doublesex* expression

**DOI:** 10.1101/2022.04.21.489080

**Authors:** Jian-jun Gao, Olga Barmina, Ammon Thompson, Bernard Kim, Anton Suvorov, Kohtaro Tanaka, Hideaki Watabe, Masanori J. Toda, Ji-Min Chen, Takehiro K. Katoh, Artyom Kopp

## Abstract

Animal evolution is characterized by frequent turnover of sexually dimorphic traits – new sex-specific characters are gained, and some ancestral sex-specific characters are lost, in many lineages. In insects, sexual differentiation is predominantly cell-autonomous and depends on the expression of the *doublesex* (*dsx*) transcription factor. In most cases, cells that transcribe *dsx* have the potential to undergo sex-specific differentiation, while those that lack *dsx* expression do not. Consistent with this mode of development, comparative research has shown that the origin of new sex-specific traits can be associated with the origin of new spatial domains of *dsx* expression. In this report, we examine the opposite situation – a secondary loss of the sex comb, a male-specific grasping structure that develops on the front legs of some drosophilid species. We show that, while the origin of the sex comb is linked to an evolutionary gain of *dsx* expression in the leg, sex comb loss in a newly identified species of *Lordiphosa* (Drosophilidae) is associated with a secondary loss of *dsx* expression. We discuss how the developmental control of sexual dimorphism affects the mechanisms by which sex-specific traits can evolve.

## Introduction

Most animals are sexually dimorphic, but the traits that distinguish males from females vary greatly from species to species. This simple observation shows that the evolution of new sex-specific traits is common in animal evolution. The origin and maintenance of sexual dimorphism can be driven by a number of evolutionary forces, most importantly by sexual selection and intragenomic conflict (Andersson, 1994; Bonduriansky and Chenoweth, 2009; Cox and Calsbeek, 2009; Darwin, 1871; van Doorn, 2009; Hill et al., 2019; Lande and Arnold, 1985; Lund-Hansen et al., 2020).

However, the evolution of sexual dimorphism is not a one-way ratchet – many ancestral sex-specific characters are lost, just as new ones are gained (Wiens, 2001). Examples of secondary losses of male-specific, sexually selected traits include the namesake tail “sword” in swordtail fish (Kang et al., 2013), dorsal sailfins in sailfin mollies (Ptacek et al., 2011), conspicuous male pigmentation in phrynosomatid lizards (Wiens, 1999) and manakins (Ribeiro et al., 2015), and vocal sacs and nuptial pads in fanged frogs (Emerson, 1996). The loss of male traits often correlates with the loss of female preferences for these traits (Morris, 1998; Morris et al., 2005; Wiens, 2001). In a swordtail species with absent or reduced male swords, females do not show any preference for longer swords, and may actually prefer swordless males or discriminate against exaggerated swords (Rosenthal et al., 2002; Wong and Rosenthal, 2006). Similarly, the secondary loss of the male sailfin in *Poecilia latipunctata* correlates with the loss of female visual preference for large-finned males (Ptacek et al., 2011). In other cases, ecological factors may result in selection against sexual dimorphism. For example, pressure from acoustically orienting parasitoid flies has led to the loss or dramatic reduction of male-specific stridulatory and resonator structures in *Teleogryllus oceanicus* crickets (Bailey et al., 2019; Pascoal et al., 2014; Zuk et al., 2006) – despite sexual selection continuing to favor stridulating males (Tanner et al., 2019).

The molecular mechanisms behind the gain and loss of sex-specific traits may depend on the developmental control of sexual dimorphism. In vertebrate animals, where the gonad non-autonomously controls sex-specific differentiation in the rest of the body by secreting male- or female-specific hormones (but see (Ioannidis et al., 2021) for an important exception), the evolution of somatic sexual traits must be mediated primarily by changes in the tissue-specific responses to hormonal signaling. In swordtails, for example, testosterone analogs induce sword development in females of sword-bearing species, but can only induce small vestigial “swordlets” or do not induce sword development at all in males of swordless species (Gordon et al., 1943; Grobstein, 1942; Sangster, 1948; Schartl et al., 2021; Zander and Dzwillo, 1969). In insects, on the other hand, sexual differentiation is predominantly cell-autonomous – that is, whether a cell will undergo male- or female-specific development depends on the expression of sex-determining genes in that cell itself (Baker and Ridge, 1980; Camara et al., 2008; Kopp, 2012; Robinett et al., 2010). This mode of sexual differentiation has important implications for the evolution of sex-specific traits.

Sex-specific cell differentiation in *Drosophila* and other insects is mostly directed by the *doublesex* (*dsx*) transcription factor, which is spliced into a female-specific isoform (*dsxF*) in females and a male-specific isoform (*dsxM*) in males (Baker et al., 1989; Burtis and Baker, 1989; McKeown, 1992; Nagoshi and Baker, 1990; Wexler et al., 2019). The two *dsx* isoforms have opposite effects on sex differentiation: *dsxM* promotes the development of male-specific traits and represses female-specific traits, while *dsxF* promotes female-specific and represses male- specific characters (Baker and Ridge, 1980; Goldman and Arbeitman, 2007; Jursnich and Burtis, 1993; Waterbury et al., 1999). Sex-specific cell differentiation is due to the distinct effects of DsxM and DsxF proteins on target gene expression (Arbeitman et al., 2016; Burtis et al., 1991; Shirangi et al., 2009; Williams et al., 2008). Importantly, *dsx* is transcribed in controlled spatial patterns (Hempel and Oliver, 2007; Kijimoto et al., 2012; Ledón-Rettig et al., 2017; Loehlin et al., 2010; Palmer and Kronforst, 2020; Rideout et al., 2010; Robinett et al., 2010; Rohner et al., 2021; Sanders and Arbeitman, 2008; Tanaka et al., 2011), and it is this fact that has a profound influence on the evolution of sexual dimorphism. For any sex-specific structure to evolve, *dsx* must be expressed in the cells that either give rise to that structure or induce it by cell-cell signaling. In tissues that already express *dsx*, sexual dimorphism can originate if Dsx acquires new downstream targets through *cis*-regulatory changes that create new, or higher affinity, Dsx binding sites (Shirangi et al., 2009; Williams and Carroll, 2009; Williams et al., 2008). If, on the other hand, the tissue is ancestrally monomorphic and does not express *dsx*, changes in the spatial regulation of *dsx* must be a necessary first step. In other words, the origin of new sex- specific traits in insects can be linked to the evolution of new *dsx* expression domains (Hopkins and Kopp, 2021; Kopp, 2012; Tanaka et al., 2011).

The best-studied example where this mechanism is at work is the origin of male-specific grasping structures on the front (T1) legs in *Drosophila* and related genera (Kopp, 2011; Rice et al., 2018; Tanaka et al., 2011). One of these, the “sex comb” found in *D. melanogaster* and its close relatives, develops from mechanosensory bristles that are typically present in both sexes. In females, these precursor bristles retain their default morphology and position. In males, however, the bristle shafts destined to become the “teeth” of the sex comb undergo several modifications, including increased length, thickness, curvature, and melanization. In many species, the entire sex comb also undergoes a coordinated 90-degree rotation, migrating from an originally transverse orientation (perpendicular to the proximo-distal axis of the leg) to a longitudinal one (parallel to the PD axis).

The sex comb is a recent evolutionary innovation that is absent in most *Drosophila* species (Kopp, 2011). It has been well studied in the Old-World *Sophophora* clade consisting of the *melanogaster* and *obscura* species groups of *Drosophila* (Fig. 1). In species that primitively lack sex combs, *dsx* is not expressed in the pupal T1 legs at the corresponding stage of development, consistent with the lack of morphological dimorphism and with the cell-autonomous model of sexual differentiation. In sex comb-bearing species, pupal *dsx* expression in the T1 leg prefigures the position and size of the sex comb, and *dsx* acts together with the HOX gene *Sex combs reduced* (*Scr*) to determine sex comb morphology (Barmina and Kopp, 2007; Kopp, 2011; Rice et al., 2019; Tanaka et al., 2011).

**Figure 1.**
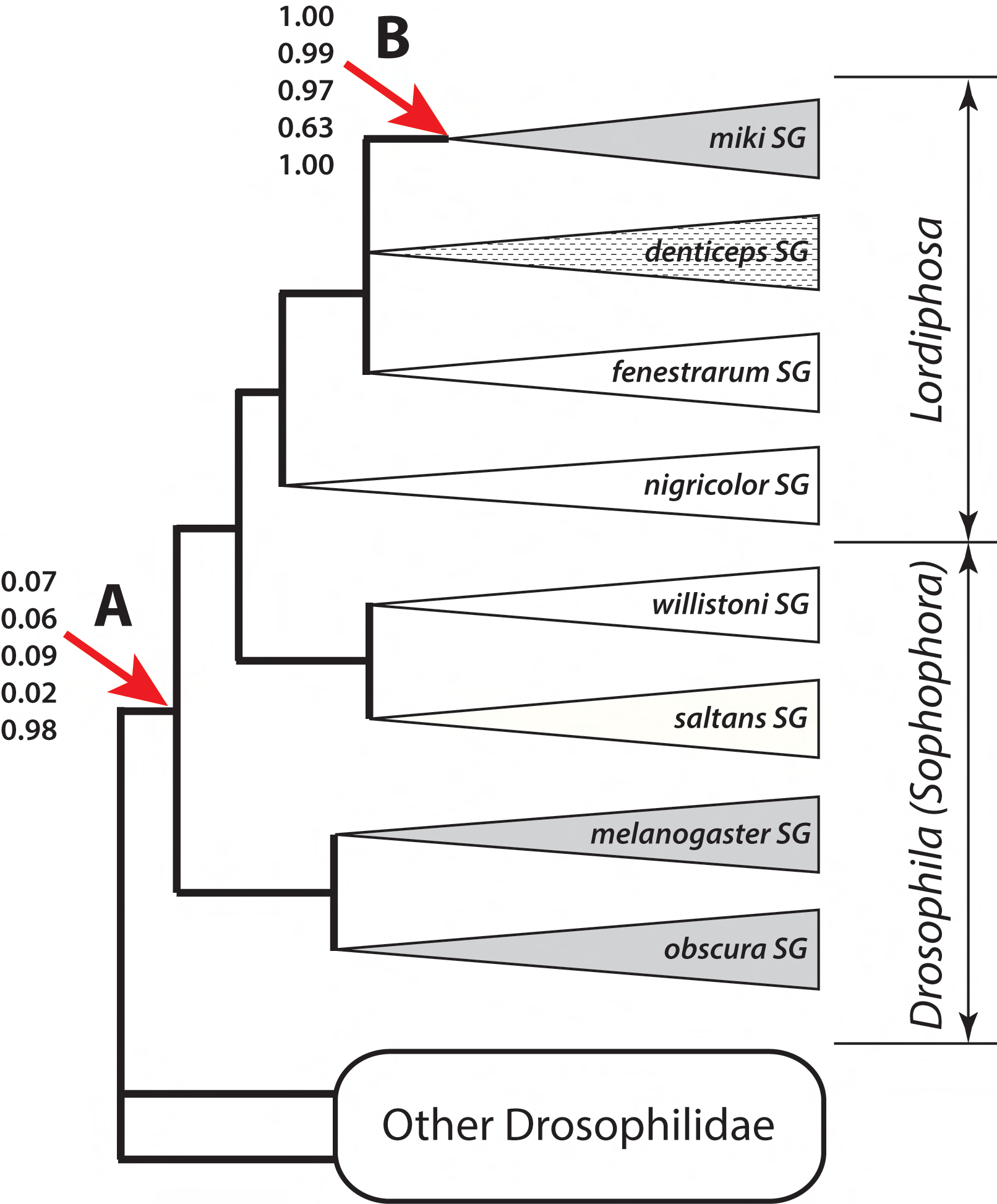
Origin and secondary loss of sex combs in *Lordiphosa*. This summary tree shows phylogenetic relationships among *Lordiphosa* (*miki, denticeps, fenestrarum*, and *nigricolor* species groups), New-World *Sophophora* (*willistoni* and *saltans* species groups), and Old-World *Sophophora* (*melanogaster* and *obscura* species groups). See Figure S4 for species-level phylogenetic relationships. The branching order of the *miki, denticeps*, and *fenestrarum* groups has not been resolved with confidence. Sex combs are present in all or almost all species of the *miki, melanogaster*, and *obscura* species groups (grey triangles), and absent in all species of the *willistoni, saltans, nigricolor*, and *fenestrarum* species groups (white triangles). Most species of the *denticeps* species group have simple sex combs (striped triangle). Numbers at nodes A (last common ancestor of *Sophophora* + *Lordiphosa*) and B (last common ancestor of the *miki* species group) denote the estimated probabilities that the last common ancestor of that clade had sex combs, under five different models of trait evolution in the following order from top to bottom: MK model with unequal rates and strict molecular clock (Fig. S5A), MK model with unequal rates and random local relaxed clock (Fig. S5B), hidden states variable rates model with two latent rate classes (Fig. S5C), modified threshold model with 9 ordered latent states (Fig. S5D), and an approximation of Dollo model (Fig. S5E). See Methods and Table S5 for model descriptions.

In addition to the Old-World *Sophophora*, sex combs are found in some species of the less well studied genus *Lordiphosa,* especially in the *miki* species group (Fartyal et al., 2017; Gao et al., 2011; Katoh et al., 2018; Kopp, 2011; Lee, 1959; Okada, 1956) (Fig. 1). All described members of this group (*L. miki*, *L. stackelbergi* (= *nipponica*), *L. magnipectinata*, and *L. clarofinis*) were at some point assigned to the subgenus *Sophophora* of the genus *Drosophila* (Bock and Wheeler, 1972; Kikkawa and Peng, 1938; Lee, 1959; Okada, 1956; Wheeler, 1949). They were subsequently transferred to the subgenus *Lordiphosa* (Lastovka and Maca, 1978; Okada, 1984), which was then elevated to the generic rank by Grimaldi (Grimaldi, 1990). Later phylogenetic analyses identified *Lordiphosa* as sister to the New-World *Sophophora* (the *willistoni* and *saltans* species groups), making *Sophophora* paraphyletic (Gao et al., 2011; Hu and Toda, 2000) (Fig. 1). This finding also raised the question whether the sex combs of Old-World *Sophophora* and *Lordiphosa* shared a common origin, or evolved independently in the two clades (Kopp, 2011). The cellular mechanism of sex comb development in *L. magnipectinata* differs from what is seen in the *melanogaster* and *obscura* species groups of *Sophophora* (Atallah et al., 2012). However, this cannot be seen as conclusive evidence of independent evolution, since the cellular mechanisms of sex comb development also differ within the *melanogaster* species group, despite the unquestionably common origin of sex combs in that lineage (Tanaka et al., 2009).

In this report, we identify a new species of *Lordiphosa* that has secondarily lost the male sex comb, making its forelegs sexually monomorphic at the morphological level. We show that this species has also lost *dsx* expression in the T1 leg. This result reinforces the close link between the evolution of sex-specific traits and *dsx* regulation in insects: just as the origin of new sexually dimorphic traits can be linked to the evolution of new *dsx* expression domains, the reversion to sexual monomorphism can be caused by tissue-specific loss of *dsx* expression.

## Materials and Methods

### 1. *Lordiphosa* collection, identification and culturing

Individuals of the *L. clarofinis* species complex, including *L. clarofinis* (Lee, 1959) and three related, undescribed species (provisionally named sp.1, sp.2, and sp.3) were collected in and near forested habitats by net sweeping over herbaceous vegetation dominated by *Galinsoga parviflora* Gav. (Fig. S1A), *Alternanthera philoxeroides* (Mart.) Griseb. or *Impatiens* spp. Collection locations (Fig. S2) are listed in Supplement Table 1. Specimens for phenotypic and phylogenetic analysis were fixed in 70% or 100% alcohol, and then identified and sexed under a stereomicroscope. In addition, isofemale lines were established from wild-caught females following (Atallah et al., 2012), maintained in glass vials (30 mm in diameter, 100 mm in height) plugged with cotton and padded with filter paper and containing a small piece of apple as adult food and shredded leaves and stems of *G. parviflora,* sterilized by freezing beforehand, as larval food (Fig. S1B-D). The founder female of each line was later subjected to molecular identification using DNA sequences of the internal transcribed spacer-1 (ITS1) region.

To distinguish the four species of the *clarofinis* complex, we relied on a combination of morphological, molecular, and geographic evidence. Old specimens of *L. clarofinis* from Korea were obtained from Dr. Nam-Woo Kim (Daegu Haany University) for help with species identification. Except for sp.3, which is known exclusively from Yunnan and is unique in lacking sex combs on male forelegs (see below), the remaining three species of the *clarofinis* complex are difficult to distinguish by external morphology. There is a slight but distinct differentiation in male genitalia between *L. clarofinis* and sp.1: ventral postgonite (ventral branch of aedeagal basal process, inner paraphysis) is distally narrow and strongly curved backward in *L. clarofinis*, but less narrow and only gently curved in sp.1 (Fig. S3). Sp.2, which is geographically restricted to the Sichuan Basin, has smaller sex combs on the first tarsomere than most populations of *L. clarofinis* (see below). Although the shape of pregonite (paramere, outer paraphysis) is variable in lateral view in *L. clarofinis*, sp.2 and sp.3, this variation appears to be due to developmental plasticity and was not used in species delimitation. However, all species including sp.2 could be readily distinguished from each other based on DNA sequences of the ITS1 region (Ji et al., 2003), and most of them could also be separated based on mitochondrial *COI* sequences (see below). *L. acongruens* (Zhang and Liang, 1992) was collected together with sp.3 in Kunming in May-June of 2018 (Supplement Table 1), cultured in the same manner as the *clarofinis* complex species, and identified by morphology.

Specimens of *L. magnipectinata* (Okada, 1956), *L*. *stackelbergi* (Duda, 1935), *L. mommai* (Takada and Okada, 1960) and *L*. *collinella* (Okada, 1968) were collected by net sweeping over undergrowth vegetation of spring ephemeral plants in Sapporo (the campus of Hokkaido University and the Jozankei area) and Iwamizawa (Hokkaido) between late May and mid-June of 2019 (Supplement Table 2). Some of the collected specimens were preserved in 100% ethanol and later used for genome sequencing. The remaining individuals, separately for each species, were kept alive in culture vials with standard *Drosophila* media in an incubator at 18 ± 0.5°C and a photoperiod of 16 h light : 8 h dark (LD 16:8) for several days. To obtain their offspring, the field-collected flies of each species were released into a plastic lidded plate (8.5 cm in diameter, 2.8 cm in height) with decayed leaves and stems of *Anemone flaccida* F. Schmidt (Ranunculaceae) and soaked filter paper at the bottom. The host plant material had been frozen at –20°C before use to kill all live insects. The female flies were then introduced to lay eggs for approximately 24 h under 18 ± 0.5°C and LD 16:8. After the adult flies were removed from the oviposition plates, the larvae were maintained under the same conditions and fed by adding decayed *A*. *flaccida* material.

### 2. Genome assemblies

The genome assemblies of *L. clarofinis, L. stackelbergi, L. magnipectinata*, *L. collinella*, and *L. mommai* are described in Kim et al. (Kim et al., 2021). We generated similar *de novo* assemblies for sp.2 and sp.3; we did not sequence the genome of sp.1 due to lack of sufficient material. For *L. clarofinis*, we sequenced the genome of a strain that was established from a single female collected in Qianling Park, Guiyang City, Guizhou Province, China, in May 2018 and inbred by single-pair, full-sib crosses for 2 generations. For sp.2, we sequenced a non-inbred isofemale line established from a female collected in Longhua Town, Pingshan County, Sichuan Province, China in May 2019. For *L. magnipectinata, L. stackelbergi, L. collinella*, and *L. mommai*, genome sequencing was performed using pools of wild-caught individuals collected in May-June 2019 in Sapporo, Hokkaido, Japan (Supplement Table 2). While it was not ideal to sequence pools of non-inbred individuals for genome assembly, we found it necessary to sequence groups of at least 20 flies to obtain sufficient material for Oxford Nanopore sequencing and resorted to computational tools to remove resulting haplotypic duplication in the assembly. For sp.3, we first established a mass culture from a small number of individuals collected in Tanglangchuan, Kunming, in May 2018. An inbred derivative of this strain (b16-1-13) was generated by three generations of single-pair, full-sib crosses, and 25 individuals from this inbred strain were used for Oxford Nanopore long-read sequencing performed by Nextomics Biosciences (Wuhan, China). For sp.2, sequencing was performed following the protocol described by Kim et al. (Kim et al., 2021). Briefly, high molecular weight genomic DNA was extracted from adult flies by phenol-chloroform extraction and prepared for Oxford Nanopore sequencing with a modified version of the SQK-LSK109 ligation kit protocol. Illumina libraries were prepared from a reserved portion of the same gDNA with a modified version of the Nextera XT Library Prep Kit protocol (Baym et al., 2015) and sequenced by Admera Health (South Plainfield, NJ) on a HiSeq 4000 machine. A detailed open-source protocol is provided at Protocols.io (dx.doi.org/10.17504/protocols.io.bdfqi3mw).

We followed the genome assembly workflow described in Kim et al. (Kim et al., 2021) to generate *de novo* assemblies, with software updated to the latest versions (as of December 2021). Nanopore reads were basecalled with Guppy v5.0.11 using the super-accuracy model. The initial read set was assembled with Flye v2.9 (Kolmogorov et al., 2019). A custom repeat library was built from this initial assembly with RepeatModeler2 (Flynn et al., 2020), then repeat masking was performed with RepeatMasker (Smit et al., 2013) using the custom library only. Duplicated sequences in the assembly were identified and removed with Purge Haplotigs (Roach et al., 2018). Regions annotated as repeats from the previous step were masked while running the Purge Haplotigs pipeline. The purged assembly was then polished with Nanopore reads, with two rounds of Racon (Vaser et al., 2017) and one round of Medaka v1.3.4. A final round of polishing using Illumina reads was performed with Pilon 1.23 (Walker et al., 2014), fixing only base-level (SNP and small indel) errors. Finally, sequences identified by a NCBI Nucleotide BLAST+ 2.12.0+ (Camacho et al., 2009) query as having a subject taxonomy ID matching non-arthropod groups were removed with BlobTools (Laetsch and Blaxter, 2017). No additional scaffolding was performed on the purged contigs.

This assembly process resulted in fairly contiguous, highly complete, and accurate assemblies. Both sp.2 and sp.3 sequencing runs had over 50% of the data in reads longer than 25 kbp. These relatively long reads resulted in final assemblies of *Lordiphosa* sp.2 of size 402.5 Mbp with a contig N50 of 217 Kbp, and *L.* sp.3 of size 432.7 Mbp with a contig N50 of 1.3 Mbp. While the lower contiguity relative to other drosophilid genome assemblies is expected due to the high genetic diversity contained in the sequenced samples, BUSCO analyses determined the gene content of the assemblies to be highly complete (sp.2: 96.8% complete; sp.3: 98.7% complete) with little duplication (sp.2: 3.8% duplicated; sp.3: 1.2% duplicated). Furthermore, the sizes of each assembly are consistent with previously sequenced *Lordiphosa* genomes and with estimates based on read depth in single-copy BUSCO genes (Supplementary Table 3). Base-level quality score analysis was performed by aligning Illumina reads back to each genome assembly with BWA (Li and Durbin, 2009), removing PCR duplicates with sambamba (Tarasov et al., 2015), and calling variants with bcftools (Li, 2011). After excluding genomic regions masked by RepeatMasker and filtering sites with base and genotype quality scores less than 30, counts of homozygous non-reference variants (that is, variants contained in the reads that do not appear in the reference) were divided by the total number of callable sites to estimate the error rate, similar to previous work (Kim et al., 2021; Solares et al., 2018). The accuracy of sp. 2 and sp. 3 assemblies was measured to be about 99.97% and 99.99% respectively (or Phred-scaled QV35 and QV39), exceeding the accuracy threshold at which most genes would be expected to have less than 1 error in their coding sequences (Koren et al., 2019). Additional details of all *Lordiphosa* assemblies are available in Supplementary Table 3.

### 3. Phylogenetic tree reconstruction

To generate a multi-locus nuclear phylogeny, we selected 250 single-copy genes identified in the genome assemblies of the seven *Lordiphosa* species and 200 other species of Drosophilidae (Supplementary Table 4). In this phylogeny, each *Lordiphosa* species was represented by a single genotype. First, we estimated multiple sequence alignments (MSAs) using the L-INS-I strategy in MAFFT v7.453 (Katoh and Standley, 2013), which generates the most accurate MSAs. Sites containing less than 3 non-gap characters were removed. After trimming, the MSA lengths ranged from 325 bp to 16,465 bp with an average length of 2,289 bp. Next, for phylogenetic inference we concatenated 250 MSAs to form a supermatrix containing 572,343 sites in total. We used this supermatrix as a single partition to estimate a maximum-likelihood (ML) tree topology in IQ-TREE v1.6.5 (Nguyen et al., 2015), specifying the GTR+I+G substitution model (Fig. S4A). To estimate the support for each node in the resulting topology, we computed three different reliability measures. We did 1,000 ultrafast bootstrap (UFBoot) replicates (Minh et al., 2013), an approximate likelihood ratio test with the nonparametric Shimodaira–Hasegawa correction (SH-aLRT), and a Bayesian-like transformation of aLRT (Anisimova et al., 2011). Additionally, to account for possible effects of ILS on tree reconstruction accuracy, we estimated a species tree topology in ASTRAL v5.14.7 using individual ML gene trees inferred for each of the 250 loci (Fig. S4B). Here individual gene trees were also estimated with IQ-TREE, again specifying the GTR+I+G substitution model. For the estimated ASTRAL tree we calculated the support of each node using local posterior probabilities (LPP) (Sayyari and Mirarab, 2016).

We implemented the Bayesian algorithm of MCMCTree v4.9h (Yang, 2007) with approximate likelihood computation to estimate divergence times using five fossils for age prior construction, analogous to the calibration scheme A described in (Suvorov et al., 2021). First, we divided our 250 loci into five equal datasets and generated five supermatrices consisting of 50 MSAs each.

We used these datasets to perform the dating analyses. For each of our five datasets, we estimated branch lengths by ML and then the gradient and Hessian matrices around these ML estimates in MCMCTree using the DNA supermatrix and species tree topology estimated by IQ-TREE. Then, we used the gradient and Hessian matrix, which constructs an approximate likelihood function by Taylor expansion (Reis and Yang, 2011), to perform fossil calibration in an MCMC framework. For this step, we specified a GTR+G substitution model with four gamma categories; birth, death and sampling parameters of 1, 1 and 0.1, respectively. To model rate variation, we used an uncorrelated relaxed clock. To ensure convergence, the analysis was run three times independently for each of the 5 datasets for 7 × 10^6^ generations (first 2 ×10^6^ generations were discarded as burn-in), logging every 1,000 generations. Additionally, we performed sampling from the prior distribution only. Convergence was assessed for the combined MCMC runs for each of the 5 datasets using Effective Sample Size criteria (ESS > 100) in Tracer (Rambaut et al., 2018).

Separately from this large taxon sample, we conducted a phylogenetic analysis of the *clarofinis* species complex where each ingroup species was represented by multiple genotypes (see Supplement Table 1 for collection locations). Two other members of the *L. miki* species group, *L. magnipectinata* and *L. stackelbergi*, were used as outgroups. DNA was extracted from each wild-caught specimen using the TIANamp^®^ Genomic DNA Kit. The mitochondrial *COI* (cytochrome oxidase subunit I) barcoding region and the nuclear ITS1 (internal transcribed spacer 1) region were amplified and sequenced using the primer pairs LCO1490/HCO2198 (Folmer et al., 1994) and CAS18sF1/CAS5p8sB1d (Ji et al., 2003), respectively. Trace files were edited in the SeqMan module of the DNAStar package 7.1.0 (DNAStar Inc., Madison, WI). The resulting sequences were aligned using the ClustalW algorithm implemented in MEGA7 (Kumar et al., 2016), and a NEXUS haplotype file was generated from the alignment in DnaSP 6 (Rozas et al., 2017). Sequences with 20 or more end gaps on either end were removed from the analysis. Neighbor-joining trees were constructed in MEGA7 from the haplotype files under the Kimura 2-parameter model, and node support was evaluated using 1000 bootstrap replicates.

### 4. Ancestral character reconstruction

To estimate the probability of sex comb origin and secondary loss, we reconstructed ancestral character states using the time-calibrated trees described above. Sex comb morphology is highly diverse, ranging from massive structures found in the *L. miki* species group (Fig. 2) and in the *ficusphila* and *montium* subgroups of the *D. melanogaster* species group to a few slightly thickened and weakly pigmented bristles that are barely distinguishable from the homologous female bristles (Kopp, 2011). The latter morphology is widespread in both *Sophophora* and *Lordiphosa*. It is found, for example, in most species of the *L. denticeps* group (Fig. 2) (Fartyal et al., 2017; Katoh et al., 2018), in some species of the *fima* subgroup (Kopp et al., 2019), and in several other species of the *D. melanogaster* group including *D. setifemur* (McEvey, 2009) and *D. ironensis*. Sex comb is also weakly developed in most species of the *ananassae* subgroup (Matsuda et al., 2009). Since our focus is on sexual dimorphism rather than any particular aspects of cell differentiation, we considered a species to have sex combs if we could detect any difference in bristle morphology (length, thickness, bluntness, and/or pigmentation) between males and females. By this permissive definition, only three species represented in our phylogenetic analysis lack sex combs entirely: *L.* sp.3, *D. prolongata*, and *D. majtoi*. This distinction is further addressed in the Discussion.

**Figure 2.**
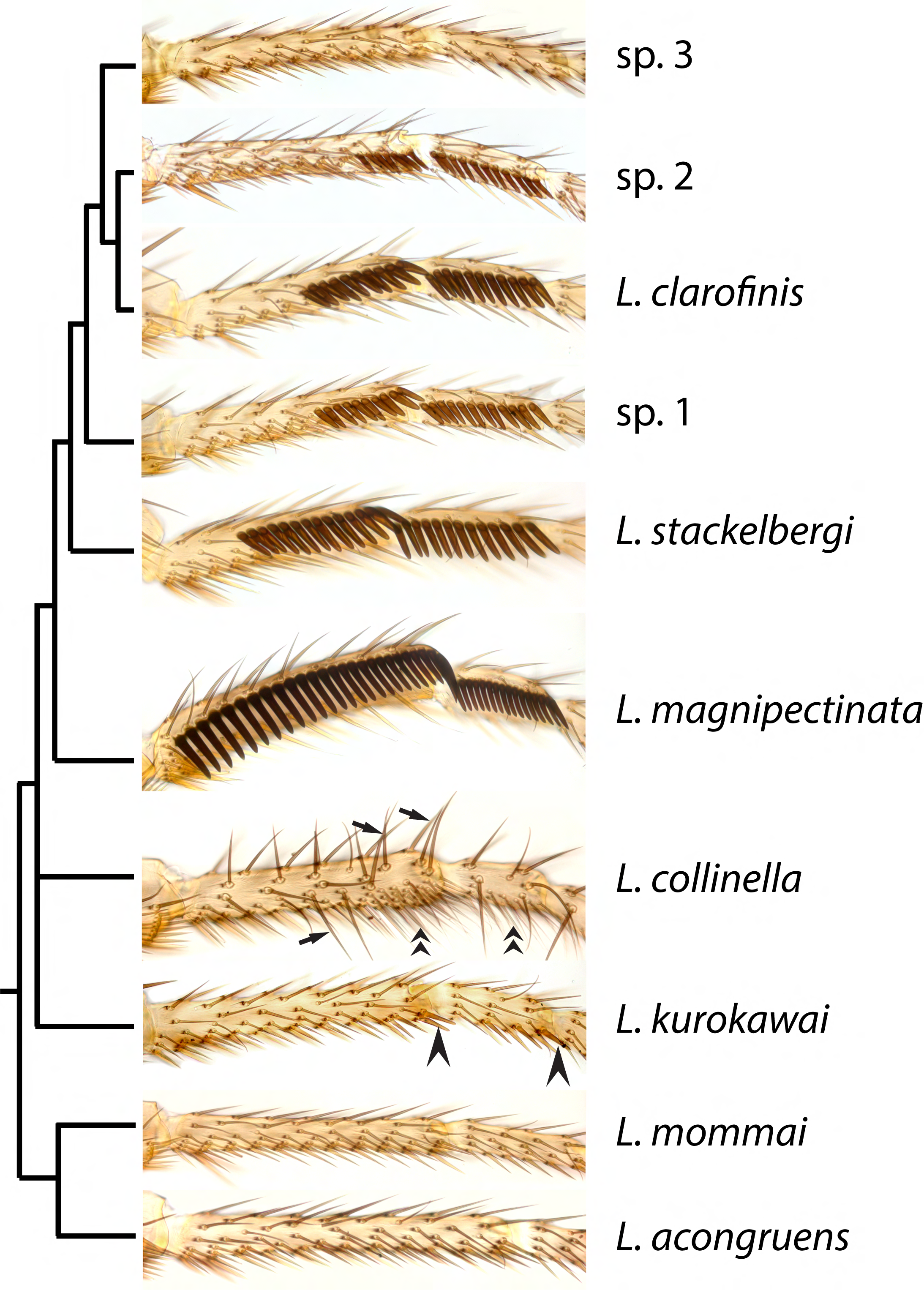
Sex combs and other male-specific leg structures in *Lordiphosa*. Males of most species of the *miki* species group, including *L. magnipectinata, L. stackelbergi, L. clarofinis*, sp.1, and sp.2, have longitudinal sex combs composed of enlarged, blunt, heavily melanized “teeth”, which are modified mechanosensory bristles. The only exception is sp.3, which lacks sex combs. Most species in the *denticeps* species group, including *L. kurokawai*, have much smaller and simpler sex combs composed of a few slightly thickened bristles (arrowheads). *L. collinella* (*fenestrarum* species group) lacks sex combs, but has other male-specific leg structures: enlarged, pointed dorsal bristles (arrows) and a ventral brush composed of thin, wavy bristles (chevrons). *L. mommai* and *L. acongruens* (*nigricolor* species group) do not show any male-specific leg modifications.

Sex comb presence/absence matrix was analyzed under five different models of trait evolution: (1) a simple MK model (Lewis, 2001) with unequal rates and a strict molecular clock; (2) an MK model with unequal rates and a random local relaxed clock (RLC; (Drummond and Suchard, 2010)); (3) a hidden states variable rates model with two latent rate classes (Beaulieu et al., 2013). The latter model assumes that each of the two binary character states (trait present vs trait absent) can exist in two discrete, not directly observable rate classes (“fast evolving / likely to change” vs ”slow evolving / not likely to change”). We also used (4) an approximation of a Dollo model made by assuming that the rate of trait loss is more than 300 times greater than the rate of gain; and (5) a modified threshold model similar to Felsenstein’s (Felsenstein, 2005). This modified threshold model assumes that the gain of a trait proceeds sequentially through 9 ordered latent states, such that each state can only transition to its nearest neighbor state towards or away from the trait. This implies that a species that lacks the trait can exist in any of the 9 “trait-absent” states that are not directly observable. For species that lack sex combs, we assume a uniform distribution over these latent states.

All models were implemented and analyzed in RevBayes (Höhna et al., 2016), and RevBayes MCMC outputs were analyzed in R using phytools (Revell, 2012). See Supplement Table 5 for prior distribution assumptions for all model parameters. Under each of the above models, except RLC, and for each of the four trees inferred from different sets of genes, MCMC was run for 50,000 cycles with a sampling rate of 1 in 50 to produce at least 200 ESS for all parameters. Because the RLC model takes significantly longer to run, we ran the MCMC for a set amount of time instead of set number of cycles. This also achieved > 200 ESS for all RLC analyses. Two independent chains were run to confirm convergence to the same posterior. Sensitivity to prior specification was assessed by comparing the marginal posterior to marginal prior for each parameter. 100 stochastic character evolution histories were simulated during one of the MCMC chains for each of the models by sampling every tenth sample from the posterior distribution. The resulting simmap files of 100 character histories mapped to a fixed tree were then analyzed to infer the ancestral states at nodes and along branches as well as the number and type of transitions. For the RLC model, we checked sensitivity to the prior on the frequency of rate shifts by comparing inferences under a prior model with expected number of rate shifts of 2 and that of 10 and obtained very similar inferences.

### 5. Adult leg imaging and phenotypic analysis

T1 legs of adult flies were dissected from the distal tibia down, mounted in Hoyers media between two coverslips, and photographed on a Leica DM5000 microscope with a Leica DC500 camera under brightfield illumination. Images were processed in Photoshop to remove the background and match exposure and color balance across species. Multiple males were examined for each species, and their sex comb phenotypes were found to be consistent within each species. We never observed any sex comb teeth in sp.3 despite examining several hundred wild-caught and lab-raised males. To examine intraspecific variation and interspecific differences in sex comb size in the remaining species of the *L. clarofinis* complex species, T1 legs of field-collected males were mounted and the numbers of sex comb teeth on the first and second tarsomeres were counted under the microscope.

### 6. Pupal leg staining

Temporary laboratory cultures of each species were established from wild-caught individuals as described above. Species identification was confirmed using a combination of morphological and molecular data. For *L. clarofinis*, we used strains collected in Zhanggou, Gaoqiao, Emeishan, Sichuan Province, China in May 2018, and in Longhua Town, Pingshan County, Sichuan in May 2019. For sp.2, the laboratory strain was also collected in Longhua in May 2019. For sp.3, we used strains collected in Kunming, Yunnan Province, China in May 2018 and in May 2019. For *L. acongruens*, we used cultured F1 progeny of flies collected in western Kunming in 2018 together with sp.3. For *L. magnipectinata, L. stackelbergi*, and *L. collinella*, the cultured strains were collected in Sapporo, Hokkaido, Japan in May 2019.

Specimens for staining were collected from lab cultures as white prepupae (0–1 hours After Puparium Formation, APF) and sexed based on the presence/absence of testes, which could be seen through the body wall. Sexed pupae were placed on a wet Kimwipe in a closed Petri dish and aged at room temperature until dissection. We observed unusually large individual variation in the rate of development within each species. Thus, instead of targeting narrow time windows, we dissected many pupae that appeared to be at the desired developmental stage based on external morphology and determined the timing of sex comb development after the fact based on the morphology of transverse bristle rows (TBRs) in the same leg. This approach allowed us to align corresponding developmental stages between species and sexes.

The dorsal half and posterior third of each pupa were cut away and remaining tissues were fixed for 30 minutes in 4% paraformaldehyde (Electron Microscopy Sciences) in 0.1 M Tris-HCl/0.3 M NaCl (pH 7.4) (TN). Fixation was followed by three washes in TN buffer. Further dissections were carried out in 0.1M Tris-HCl/0.3M NaCl (pH 7.4) containing 0.5% Triton X-100 (TNT). Legs were removed from the pupal cuticle by first tearing the cuticle around the dorsal femur/tibia boundary and then pulling out the tibial and tarsal segments. Tissues were then treated with Image-iT FX Signal Enhancer (Invitrogen) for 30min and washed three times for 15 min each in TNT. Samples were incubated with the primary antibodies overnight at 4°C. The primary antibodies used were monoclonal mouse anti-Dsx (Mellert et al., 2012) (1:5 dilution in TNT) and rat-Elav-7E8A10 (O’Neill et al., 1994) (1:10 dilution in TNT), both obtained from the Developmental Studies Hybridoma Bank. Legs were then washed in TNT buffer four times and incubated in the secondary antibodies overnight at 4°C. The secondary antibodies were Invitrogen/Thermo Fisher 488 mouse # A32723 preabsorbed against rat and Invitrogen/Thermo Fisher 555 rat #A21434 preabsorbed against mouse (1:200 dilution in TNT). Stained samples were washed four times in TNT and mounted in Fluoromount 50 (Southern Biotech). Fluorescent images were collected on an Olympus FV1000 confocal microscope at UC Davis or an Olympus FV1000 confocal microscope at the Research Institute of Resource Insects, Chinese Academy of Forestry, Kunming. The 40X lens was used, with the gain adjusted for the dynamic range of each sample. For *L. clarofinis*, sp.3, and *L. acongruens*, we imaged at least 10 males per species; fewer individuals were examined for sp.1 and sp.2. To assemble leg images, signals from non-epidermal tissues were removed from each confocal section, and Z-series projection was produced using ImageJ.

## Results

### 1. Sex comb evolution in *Lordiphosa*

Sex combs are present in almost all Old-World *Sophophora* (the *D. melanogaster* and *D. obscura* species groups) but are absent in all New-World *Sophophora* (the *willistoni* and *saltans* groups) (Fig. 1). The *Lordiphosa* genus is the sister lineage to the New-World *Sophophora* (Gao et al., 2011; Kim et al., 2021; Suvorov et al., 2022). Within *Lordiphosa*, sex combs are found in all described members of the *miki* species group and all members of the *denticeps* species group except *L. medogensis* Katoh and Gao, 2018 (Fartyal et al., 2017; Gao et al., 2011; Katoh et al., 2018) (Fig. 1). Although the phylogeny of *Lordiphosa* is not fully resolved, neither the *miki* nor the *denticeps* species groups are basal within that genus. Thus, it is possible that sex combs evolved independently two or three times: once in the Old-World *Sophophora*, and either once or twice in *Lordiphosa*. Independent origin of sex combs in *Sophophora* and *Lordiphosa* would be consistent with the differences in cellular mechanisms that underlie sex comb development in the *L. miki* species group compared to *Sophophora* (Atallah et al., 2012; Tanaka et al., 2009). However, it is also possible that sex combs evolved once at the base of the (*Sophophora* + *Lordiphosa*) clade, and subsequently underwent many independent losses: in the New-World *Sophophora*, in the *L. nigricolor* species group, in the *L. fenestrarum* species group, and in a few isolated species of the *D. melanogaster* and *L. denticeps* species groups (Fartyal et al., 2017; Kopp, 2011). To distinguish between these scenarios, we used a whole-genome dataset consisting of 250 single-copy BUSCO genes (572,343 total sites) to reconstruct the phylogeny of >200 drosophilid species and estimate the probability of the single-gain and multiple-gains models of sex comb evolution.

Under most models of character evolution, our analysis suggests that sex combs originated independently in *Lordiphosa* and in Old-World *Sophophora* (Fig. S5 A-D). The probability that the last common ancestor of both clades had sex combs is 0.09 under the hidden states variable rates model (Beaulieu et al., 2013), and 0.02-0.07 under the MK models (Drummond and Suchard, 2010; Lewis, 2001) and the threshold model with latent ordered states (Fig. 1). As expected, enforcing an approximate Dollo model by assuming a highly informative prior where the rate of trait loss is more than 300 times higher than the rate of trait gain leads to a different conclusion, namely that the sex comb originated only once at the base of all Drosophilidae and was subsequently lost many times (Fig. S5 E). This result may be partly driven by biased taxon sampling, since the Old-World *Sophophora* are heavily overrepresented in the available set of genomes and therefore in the BUSCO phylogeny (Kim et al., 2021; Suvorov et al., 2022). However, unless we assume this extreme model of trait evolution, our analysis supports independent origin of sex combs in *Lordiphosa*, separately from the Old-World *Sophophora* (Fig. 1, Fig. S5 A-D).

### 2. *dsx* expression associated with sex comb development in *Lordiphosa*

All species of the *nigricolor* species group, which represents the most basal lineage within *Lordiphosa* (Gao et al., 2011; Kim et al., 2021), lack sex combs. We examined leg development and *dsx* expression in *L. acongruens,* a representative of this species group (Fig. 1, 2). We found that, at the stage when sex combs develop in other species of *Lordiphosa* and in Old-World *Sophophora*, Dsx expression is absent in both males and females of *L. acongruens* (Fig. 3A). This is consistent with the lack of Dsx expression at this stage in the New-World *Sophophora*, which also lack sex combs (Rice et al., 2019; Tanaka et al., 2011), as well as with the absence of sex-specific chemosensory bristles in *Lordiphosa* (Kopp and Barmina 2022). Dsx expression is also absent at earlier, prepupal stages in *L. acongruens* (Fig. 3H).

**Figure 3.**
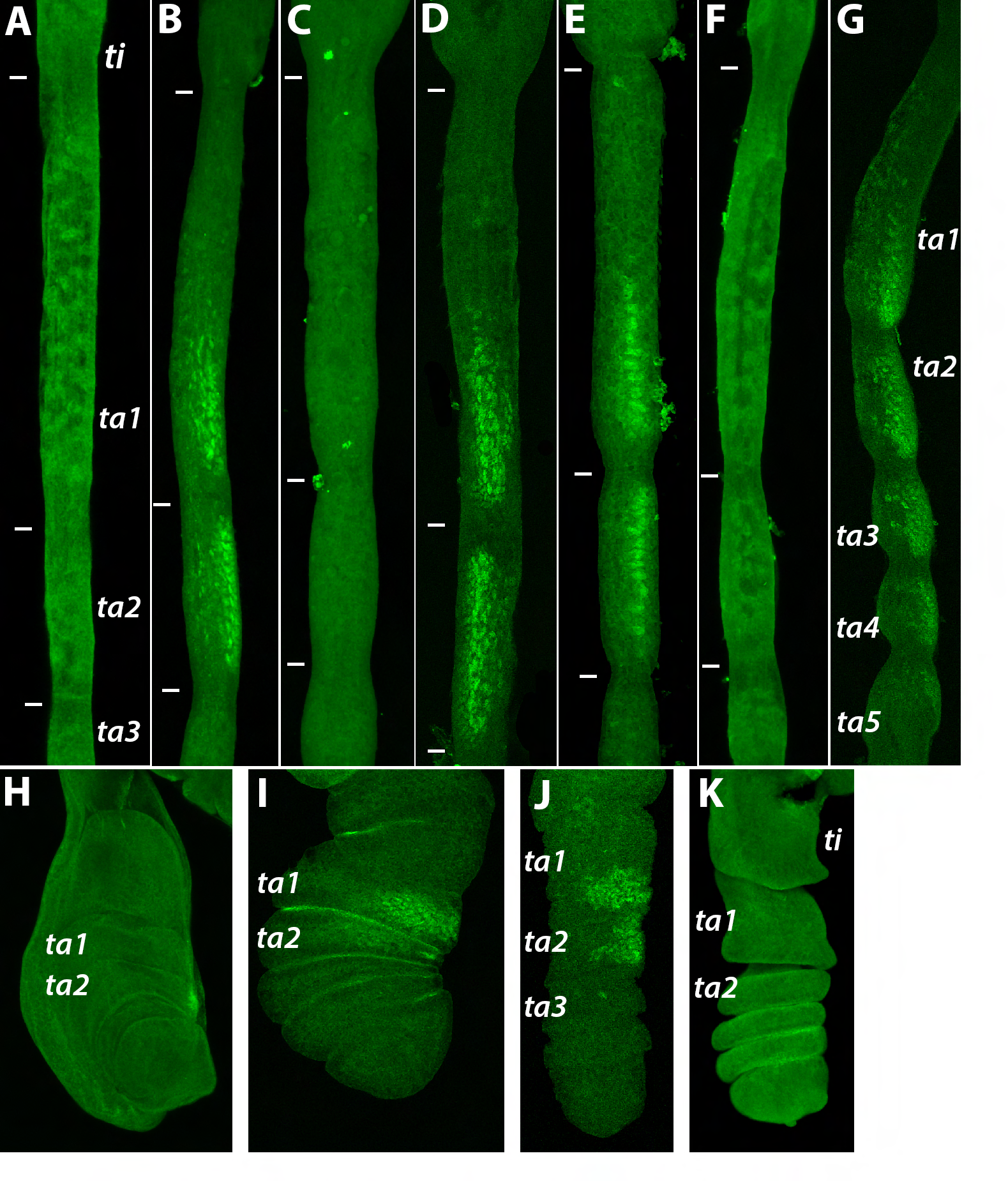
Dsx expression in *Lordiphosa* T1 legs. Proximal is up, distal is down. Tibia (*ti*) and tarsal segments 1–5 (*ta1*–*ta5*) are indicated in some panels; in other panels, the boundaries between these segments are shown with white tick marks. Panels A– G show pupal legs at stages equivalent to 24–30 hours After Puparium Formation (APF) in *D. melanogaster*. **A**. *L. acongruens* male; Dsx protein is absent. **B**. *L. clarofinis* male; Dsx is expressed on the anterior-ventral leg surface in the distal part of ta1 and all of ta2. **C**. *L. clarofinis* female; no Dsx expression is seen at this stage. **D**. *L.* sp.1 male; Dsx expression is similar to *L. clarofinis*. **E**. *L.* sp.2 male; Dsx expression is similar to *L. clarofinis*. **F**. *L.* sp.3 male; no Dsx expression is observed. **G**. *L. collinella* male; Dsx staining is present in all tarsal segments. **H**. The T1 leg imaginal disc of the white prepupa (0 hrs APF) of *L. acongruens* male; no Dsx expression is observed. **I**. Prepupal leg of *L.* sp.2 male at the stage equivalent to ∼2–3 hrs APF in *D. melanogaster*; Dsx is expressed on the anterior-ventral surface of ta1 and ta2. **J**. Prepupal leg of *L.* sp.2 male at the stage equivalent to ∼6–7 hrs APF in *D. melanogaster*. **K**. Prepupal leg of *L.* sp.3 male at the stage equivalent to ∼5–6 hrs APF in *D. melanogaster*; no Dsx expression is seen.

All species of the *L. miki* species group, except sp.3, have well-developed sex combs reminiscent of the largest sex combs seen in the *melanogaster* species group of *Sophophora* (Atallah et al., 2012; Kopp, 2011) (Fig. 2). We observed Dsx expression in the presumptive sex comb region of the prepupal and pupal T1 legs of *L. clarofinis*, sp.1, and sp.2 (Fig. 3 B, D, E, I, J). As in the Old-World *Sophophora* (Tanaka et al., 2011), the spatial domain of Dsx expression corresponds to sex comb position. Also as in *Sophophora*, Dsx expression at the pupal stage is strong in males but absent in females (Fig. 3 B, C). Thus, despite the apparently independent evolution of sex combs and the differences in the cellular mechanisms of their development in *Lordiphosa* vs *Sophophora* (Atallah et al., 2012), in both cases their origin is associated with a novel spatial domain of *dsx* expression.

Sex combs are absent in the *L. fenestrarum* species group. However, *L. collinella*, a member of this group, has different sexually dimorphic traits on the T1 leg. In this species, a sparse ventral brush and a set of enlarged dorsal spines, both derived from mechanosensory bristles, are present in males but not in females (Fig. 2). As with the sex comb, we find that the presence of these structures in adult males is prefigured by male-specific expression of Dsx at the pupal stage (Fig. 3G). This consistent association of novel sex-specific traits with newly evolved domains of *dsx* expression is predicted by the cell-autonomous model of sexual differentiation in insects (Hopkins and Kopp, 2021; Kopp, 2011; Ledón-Rettig et al., 2017).

### 3. Secondary loss of the sex comb in the *clarofinis* species complex

*Lordiphosa clarofinis* was originally described from type specimens collected in Kongju, South Korea (Lee, 1959). Our field surveys in China and Japan during the last two decades identified a much larger geographical range for *L. clarofinis* than previously known, covering the central, southern, and southwestern China except for Yunnan (Table S1 and Fig. S2). These surveys also revealed the presence of three new forms closely resembling *L. clarofinis*. *L.* sp.1 aff. *clarofinis* is found in Japan and characterized by ventral postgonite that is distally broader and has a gentler backward curvature than in *L. clarofinis* (Fig. S3)*. L.* sp.2 aff. *clarofinis* is restricted to the Sichuan Basin, and *L.* sp.3 aff. *clarofinis* occurs exclusively in Yunnan. The combination of morphological characters and molecular evidence (Fig. S6) suggests that *L. clarofinis*, sp.2, and sp.3 are distinct, but we did not test whether they are reproductively isolated from each other.

*L.* sp.3 aff. *clarofinis* is unique in the *miki* species group in completely lacking sex combs on both the first and the second tarsomeres (Fig. 2). The other three species of the *clarofinis* complex have sex combs on both tarsomeres but differ in the number of sex comb teeth, especially on the first tarsomere. The sex combs of *L. clarofinis* and sp.1 are similar in size, and both are significantly larger than in sp.2 (Fig. S7). An exception is found in the Sichuan population of *L. clarofinis*, which shows sex combs that are similar in size to sp.2, which is sympatric with this population, rather than to 10 other populations of *L. clarofinis*, which are allopatric with sp.2 (Fig. S7). The reason for the sympatric convergence between *L. clarofinis* and sp.2 is unknown.

We used single-copy BUSCO genes to reconstruct the phylogeny of the *miki* species group (the clade represented in our analysis by the *clarofinis* species complex, *L. stackelbergi*, and *L. magnipectinata*). First, a neighbor-joining tree was constructed separately from each of 10 different loci, using *L. mommai* (*nigricolor* species group) as outgroup. All 10 genes supported the ((*L. clarofinis*, sp.2) sp.3) tree topology (genome data are not available for sp.1). Nine loci supported (*magnipectinata* (*stackelbergi* (*clarofinis* complex))) topology; one supported ((*magnipectinata*, *stackelbergi*) (*clarofinis* complex)) topology. The combined 250-locus BUSCO dataset (see above) produced a credible set of only one tree: root-(*magnipectinata* (*stackelbergi* (sp.3 (*clarofinis,* sp.2)))) with 100% bootstrap and aLRT support at each node (Fig. S4A); the same topology is also observed in the ASTRAL tree (Fig. S4B).

The key question for our analysis is whether the sex comb was lost secondarily in sp.3 aff*. clarofinis*. To address this question, we estimated the probability that the last common ancestor of the *Lordiphosa miki* species group had sex combs. We found this probability to be 0.99-1.00 under the Dollo and MK models (Drummond and Suchard, 2010; Lewis, 2001) and 0.97 under the hidden states variable rates model (Beaulieu et al., 2013) (Fig. 1). An outlier result in this analysis was produced by the threshold model with latent ordered states, which accommodates more gradual transitions between the “present” and “absent” states of discrete characters. However, even this model puts the probability of secondary loss at 0.63 (Fig. 1). Thus, we conclude that the sex comb was present in the last common ancestor of the *clarofinis* species complex, but was lost secondarily in sp.3 after its divergence from the common ancestor of *L. clarofinis* and sp.2.

Separately, we constructed neighbor-joining trees of 140 mitochondrial *COI* haplotypes isolated from 650 individuals of six *Lordiphosa* species, and 36 nuclear ITS1 haplotypes sampled from 653 individuals (Table S1). Both analyses lend strong support to the monophyly of the *L. clarofinis* species complex with respect to the outgroups (*L. magnipectinata* and *L. stackelbergi*) (Fig. S6).

*L.* sp.1 aff. *clarofinis* is placed as most basal within the complex, although with weak support. The relationship among the remaining three species is not fully resolved. In the *COI* tree, *L. clarofinis* is well supported as monophyletic, but sequences from *L.* sp.2 aff. *clarofinis* and *L.* sp.3 aff. *clarofinis* form an intermingled clade, with neither species appearing as monophyletic and some haplotypes shared between sp.2 and sp.3 (Fig. S6 A). In the ITS1 tree, each of the four focal species is monophyletic. However, sp.2 and sp.3 are placed as sister taxa to the exclusion of *L*. *clarofinis* (Fig. S6 B), in contrast to the multilocus BUSCO phylogeny that places *L. clarofinis* as sister to sp.2 (Fig. S4 A, B). The incongruence between ITS1 and BUSCO tree topologies could be due to ancestral lineage sorting, post-speciation gene flow, or simply to the small amount of data in the ITS1 dataset. The *COI* gene tree is most consistent with hybridization between sp.2 and sp.3 resulting in interspecific introgression of mitochondria, which is known to occur in *Drosophila* (Bachtrog et al., 2006; Llopart et al., 2014; Nunes et al., 2010; Turelli et al., 2018). If such hybridization has indeed occurred, it could also have affected some nuclear loci, including genes that control sex comb development. However, the tree reconstructed from the ∼20 kb *dsx* non-coding region supported the same relationships as the BUSCO tree: root-(*magnipectinata* (*stackelbergi* (sp.3 (*clarofinis*, sp.2)))). Most importantly for this study, sp.3 occupies a derived position within the *miki* species group and the *clarofinis* complex in all analyses, confirming a secondary loss of the sex comb in this species.

### 4. Secondary loss of *dsx* expression in sp.3 correlates with loss of the sex comb

In both males and females of sp.3, no Dsx expression is seen in the T1 legs at the pupal stages when sex combs are developing in *L. clarofinis* and sp.2 (Fig. 3F). In most *Drosophila* species, including some that lack sex combs, *dsx* is also expressed in the T1 legs earlier, in late third instar larvae and prepupae (Tanaka et al., 2011). This early expression is necessary for the development of sex-specific chemosensory bristles as well as the sex comb (Rice et al., 2019). In sp.3., however, no Dsx expression is seen at the prepupal stage when chemosensory bristles are specified (Fig. 3K). This is consistent with the absence of sex-specific chemosensory bristles in all examined species of *Lordiphosa*, which stands in contrast with the majority of *Sophophora* and *Drosophila* species (Kopp and Barmina, 2022). We conclude that the secondary loss of the sex comb in sp.3 coincides with the loss of *dsx* expression in the presumptive sex comb region. In addition, the secondary loss of sex-specific chemosensory bristles observed in *Lordiphosa* may also correlate with the loss of the early phase of *dsx* expression in prepupal legs.

## Discussion

Our results show that the relationship between *dsx* expression and the evolution of sexual dimorphism goes both ways: while the origin of novel sex-specific traits is sometimes linked to new domains of *dsx* expression, the loss of sexually dimorphic structures can be associated with the loss of *dsx* expression. This linkage is a logical consequence of the mainly cell-autonomous control of sexual differentiation in insects: with rare exceptions, cells that transcribe *dsx* have the potential to differentiate in sex-specific ways, while those that lack *dsx* expression do not (Camara et al., 2008; Hopkins and Kopp, 2021; Kopp, 2012; Ledón-Rettig et al., 2017). It will be interesting to see whether the pattern of gains and losses of *dsx* expression extends to other models where sex-specific traits have been both gained and lost – such as beetle horns (Emlen et al., 2005; Moczek et al., 2006), Lepidopteran androconia (Prakash and Monteiro, 2020; Simmons et al., 2012; Valencia-Montoya et al., 2021), or Batesian mimicry in swallowtail butterflies (Kunte, 2009; Palmer and Kronforst, 2020).

However, the gain and loss of *dsx* expression is not the only mechanism by which sex-specific traits can be gained and lost. In many cases, *dsx* expression in the tissue can be conserved, and the tissue can retain the potential for sexually dimorphic development, even as particular sex-specific characters are gained and lost. In such scenarios, the gain and loss of sex-specific traits is caused by evolutionary changes in the response of downstream target genes to *dsx*, rather than to changes at the *dsx* locus itself. This can happen through changes in Dsx binding sites in the regulatory regions of those downstream targets – as, for example, in the evolution of sexually dimorphic abdominal pigmentation (Gompel and Carroll, 2003; Kopp et al., 2000; Rogers et al., 2013; Williams et al., 2008), or in the synthesis of sex-specific pheromones in oenocyte cells (Shirangi et al., 2009). In fact, this mechanism may operate in the evolution of sex combs, as well. In *D. prolongata* (*melanogaster* species group), the sex comb was lost, but *dsx* continues to be expressed and controls the development of sex-specific chemosensory bristles in the T1 leg (Luecke et al., 2022). Although complete secondary loss of sex combs (defined in this study as the absence of any detectable sexual dimorphism) is relatively rare, extreme reductions of the sex comb are more common. In the *D. melanogaster* species group alone, various degrees of reduction are seen in *D. setifemur, D. ironensis, D. flavohirta*, and some species of the *ananassae*, *fima*, and *takahashii* subgroups (Grimaldi et al., 2015; Kopp, 2011; Kopp et al., 2019; Matsuda et al., 2009; McEvey, 2009). Since these species still retain some level of sexual dimorphism, we expect that their attenuated sex comb morphology results from changes in *dsx* target genes rather than a loss of *dsx* expression.

Finally, the development of sexually dimorphic structures is not always cell-autonomous. For example, the development of the male-specific Muscle of Lawrence (MOL) in *Drosophila* depends not on the sex of the myoblasts that give rise to that muscle, but rather on the sex of the motoneuron that innervates it, and specifically on the expression of the *fruitless* (*fru*) gene in that neuron (Lawrence and Johnston, 1986; Nojima et al., 2010; Usui-Aoki et al., 2000). MOL has also undergone many gains and losses in *Drosophila* evolution (Gailey et al., 1997; Liang et al., 2021), which could be caused by changes either in the differentiation of the inducing motoneuron or in the response of the developing muscle to its synaptic output. Unsurprisingly, the genetic basis of trait evolution will be biased by the mode of its development.

## Supplementary figures

**Figure S1.**
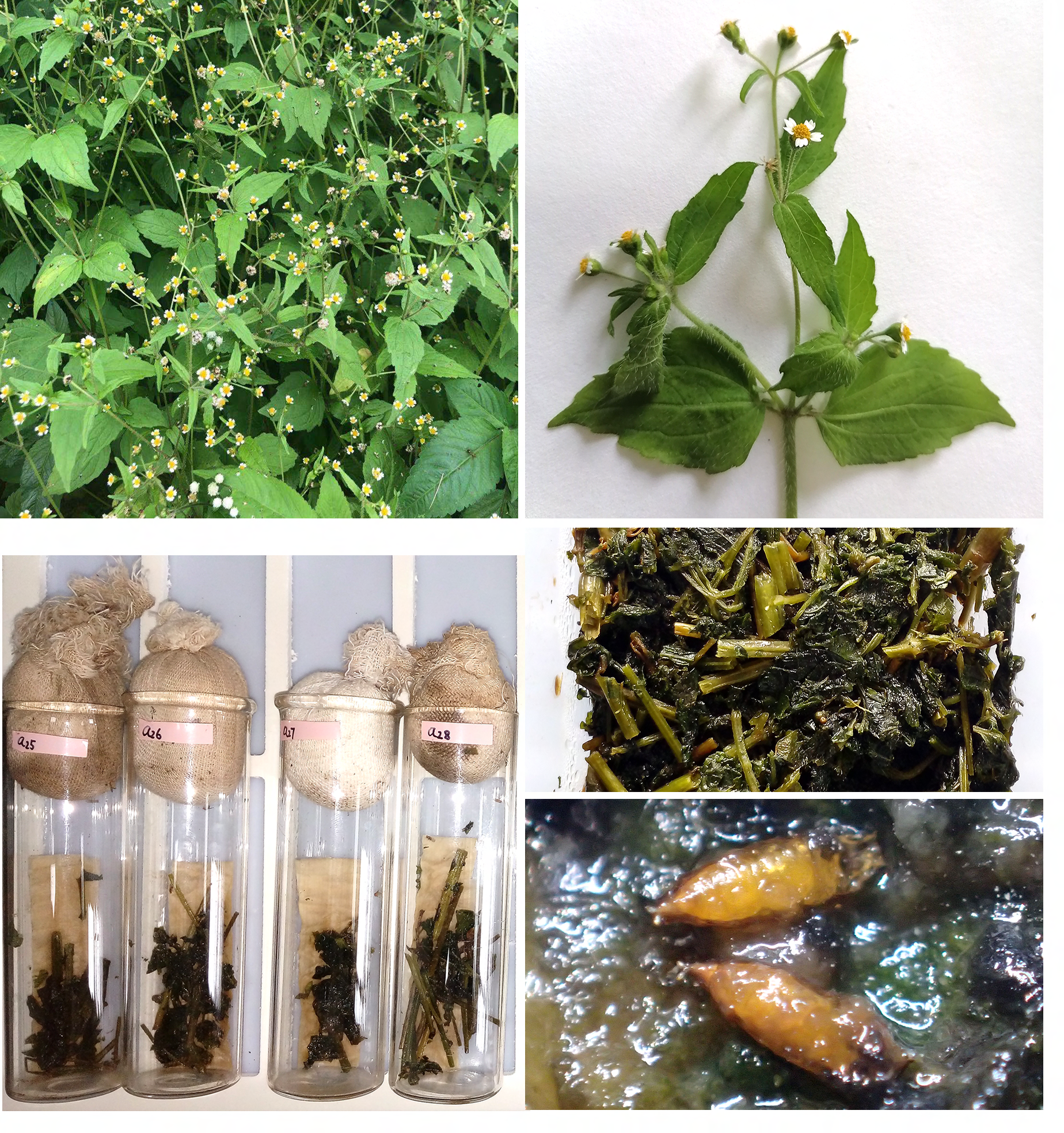
Culturing species of the *Lordiphosa clarofinis* species complex. (A) Thick growth of *Galinsoga parviflora* Gav., one of the natural breeding substrates (Baihualing, Baoshan, Yunnan). (B) Leaves and flowers of the same plant. (C) Frozen leaves and stems of *G. parviflora* used for rearing *Lordiphosa* flies. (D) Glass vials containing *G. parviflora* as larval food, and filter paper as pupation substrate (pieces of apple, used as adult food, were added later). (E) Pupae of *L. clarofinis* reared in the lab on this substrate.

**Figure S2.**
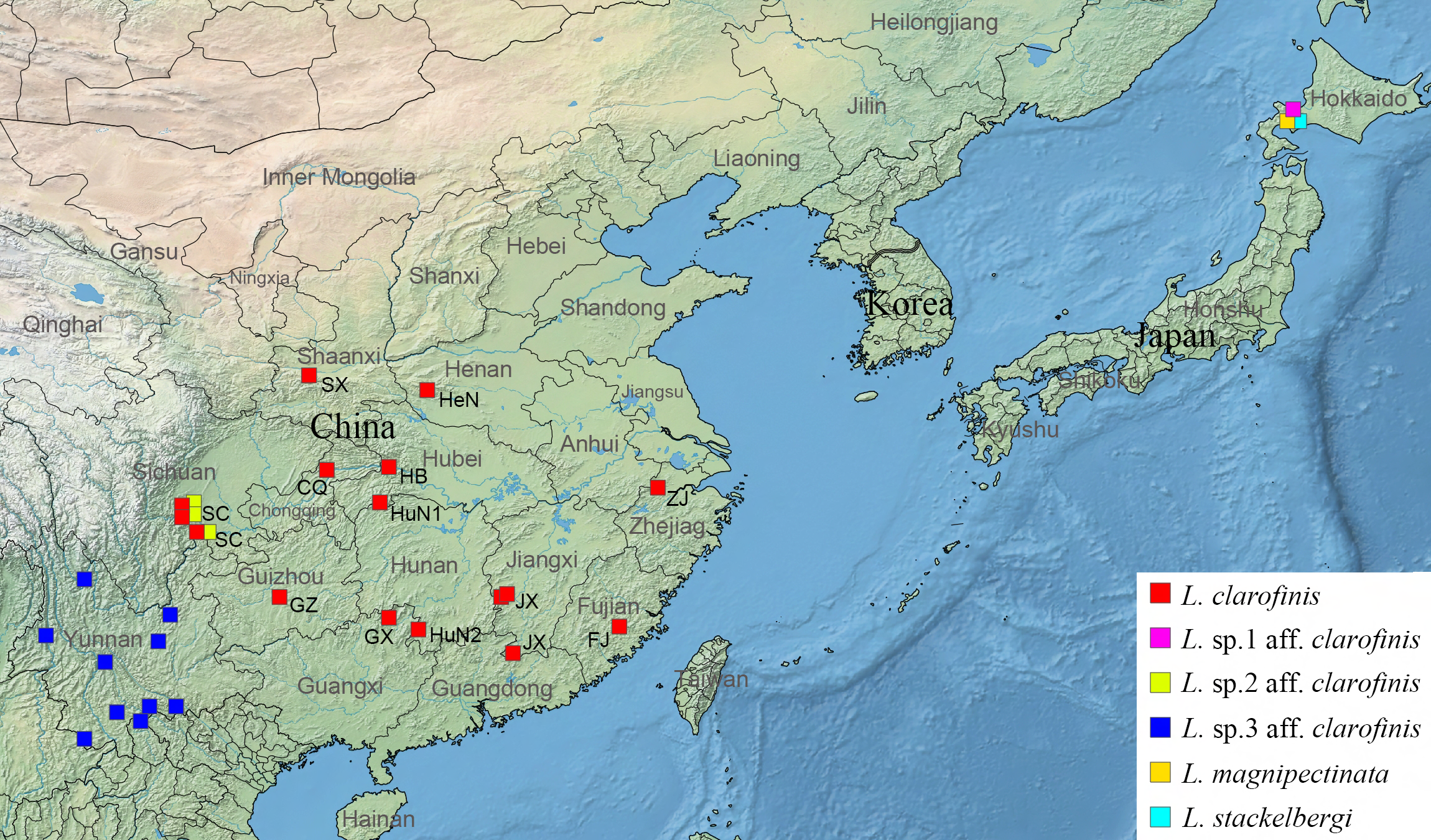
Collection sites of four members of the *L. clarofinis* complex and two outgroup species. Map was created using Simplemappr (http://www.simplemappr.net/).

**Figure S3.**
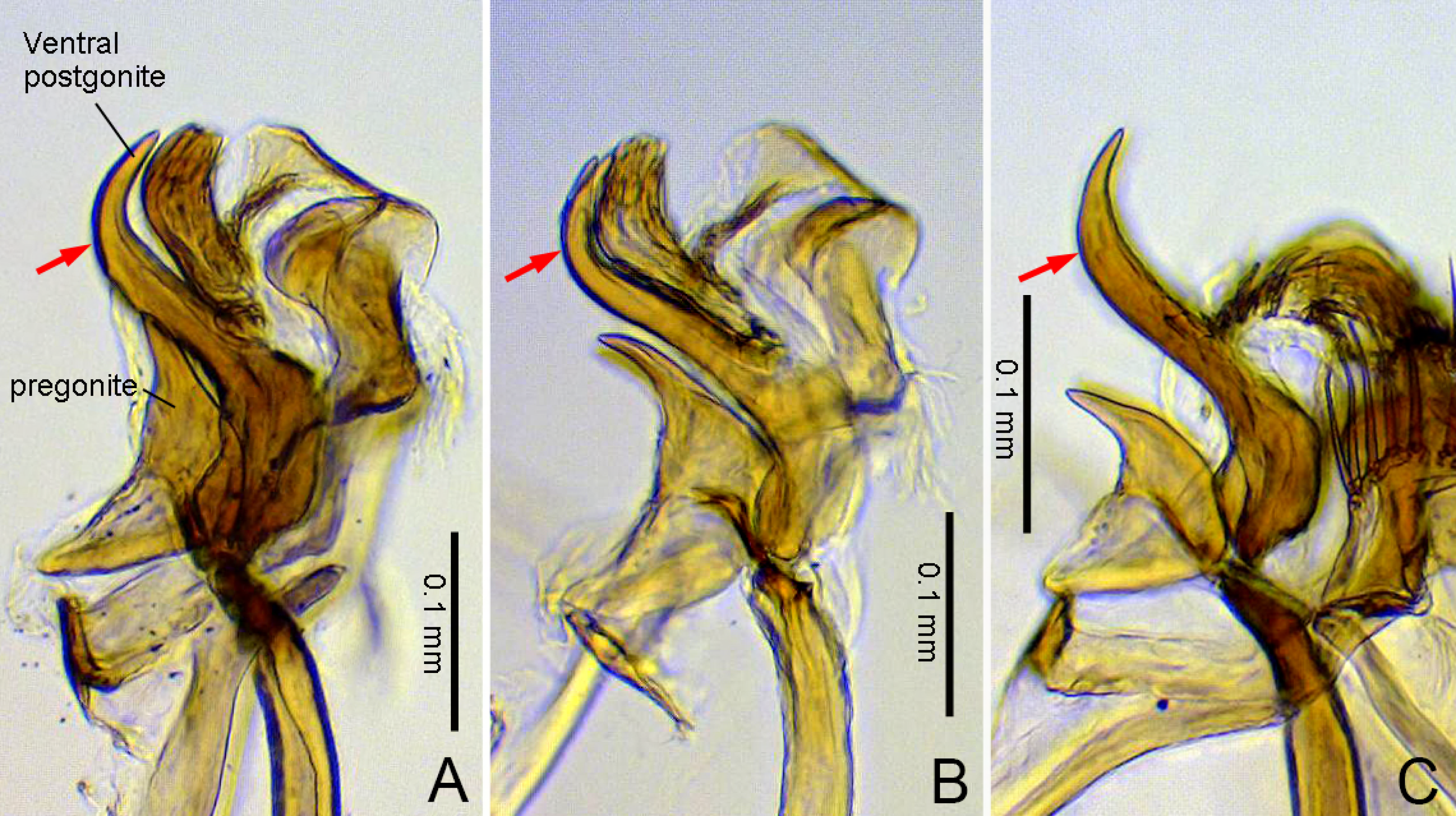
Pregonite and ventral postgonite of *L. clarofinis* (all lateral views; A, specimen from Korea; B, specimen #01711 from Taibaishan, Shaanxi, China) and *L.* sp.1 aff. *clarofinis* (C, specimen #01903 from Hokkaido, Japan). Red arrows indicate the bending position of the ventral postgonite in each specimen.

**Figure S4.**
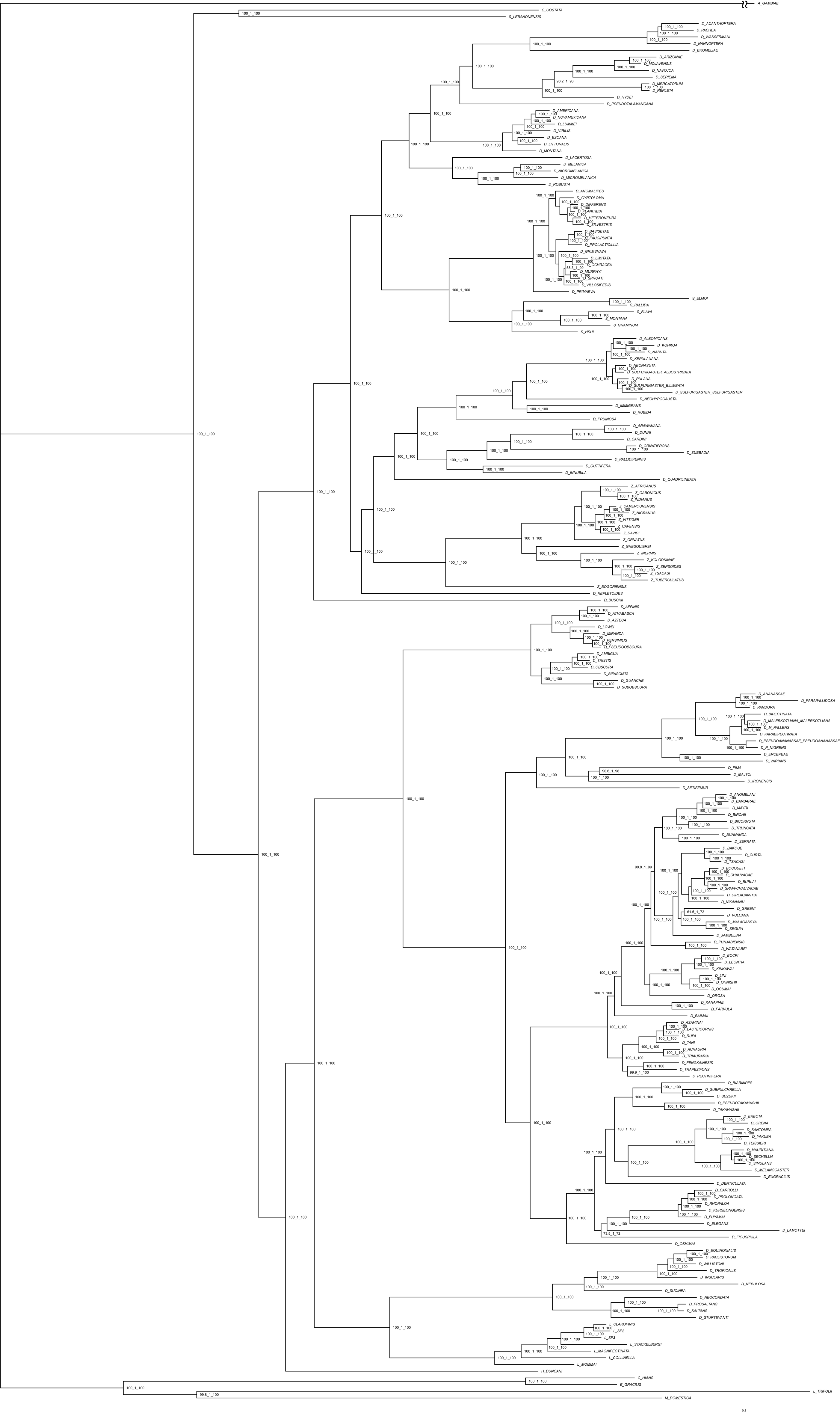
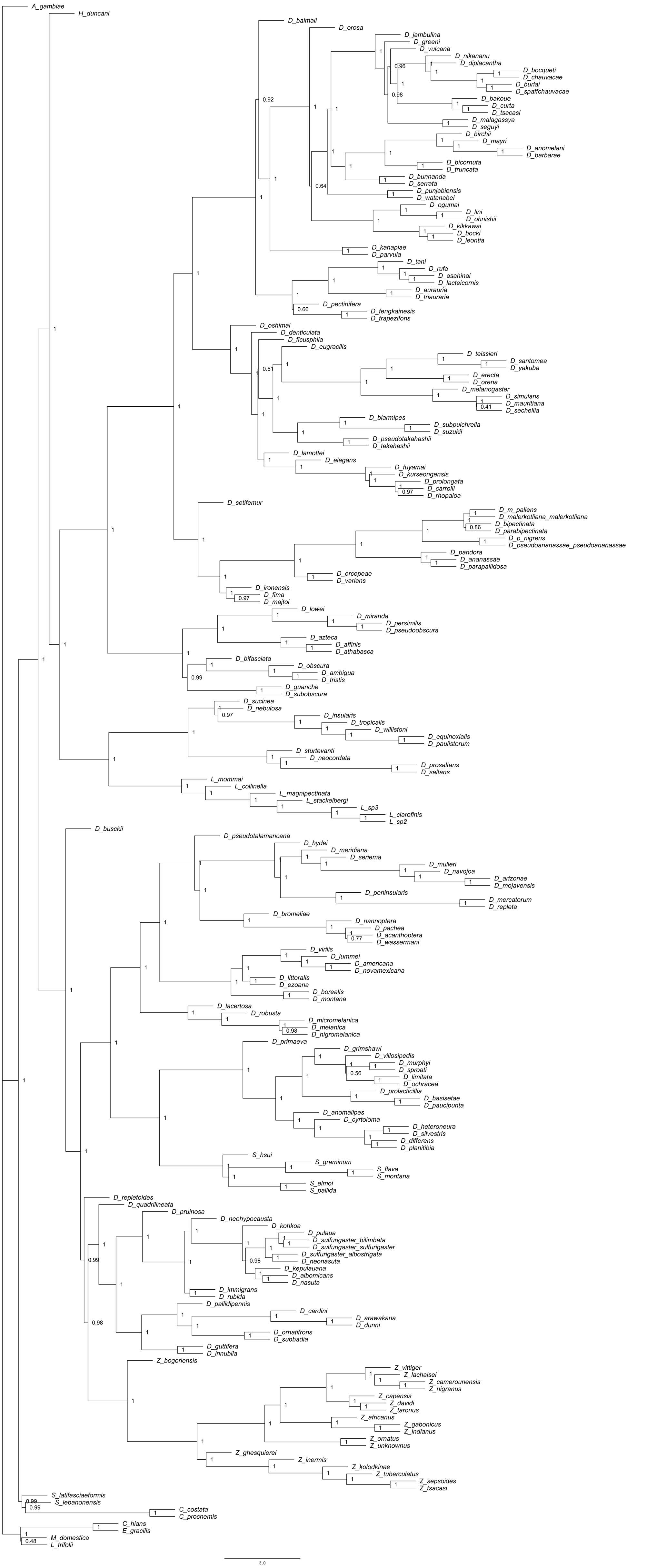
**A.** A maximum likelihood tree of 207 species of *Drosophila*, *Lordiphosa*, and related genera reconstructed using IQ-TREE v1.6.5 from 250 single-copy BUSCO loci (572,343 total sites) (Table S4) and rooted with *Anopheles gambiae*. At each node, the three measures of node support are, in order: 1000 replicates of ultrafast bootstrap (Minh et al., 2013), a Bayesian-like transformation of approximate likelihood ratio test (aLRT) (Anisimova et al., 2011), and aLRT with the nonparametric Shimodaira–Hasegawa correction (SH-aLRT). The *A. gambiae* branch is not to scale. See Methods for details of phylogeny reconstruction. **B.** ASTRAL tree topology, with local posterior probability support measures for each node, estimated from the 250 individual gene trees. Branch lengths are reported in coalescent time units. ASTRAL only reports the length of internal branches; the length of each terminal branch was manually set to 1.

**Figure S5.**
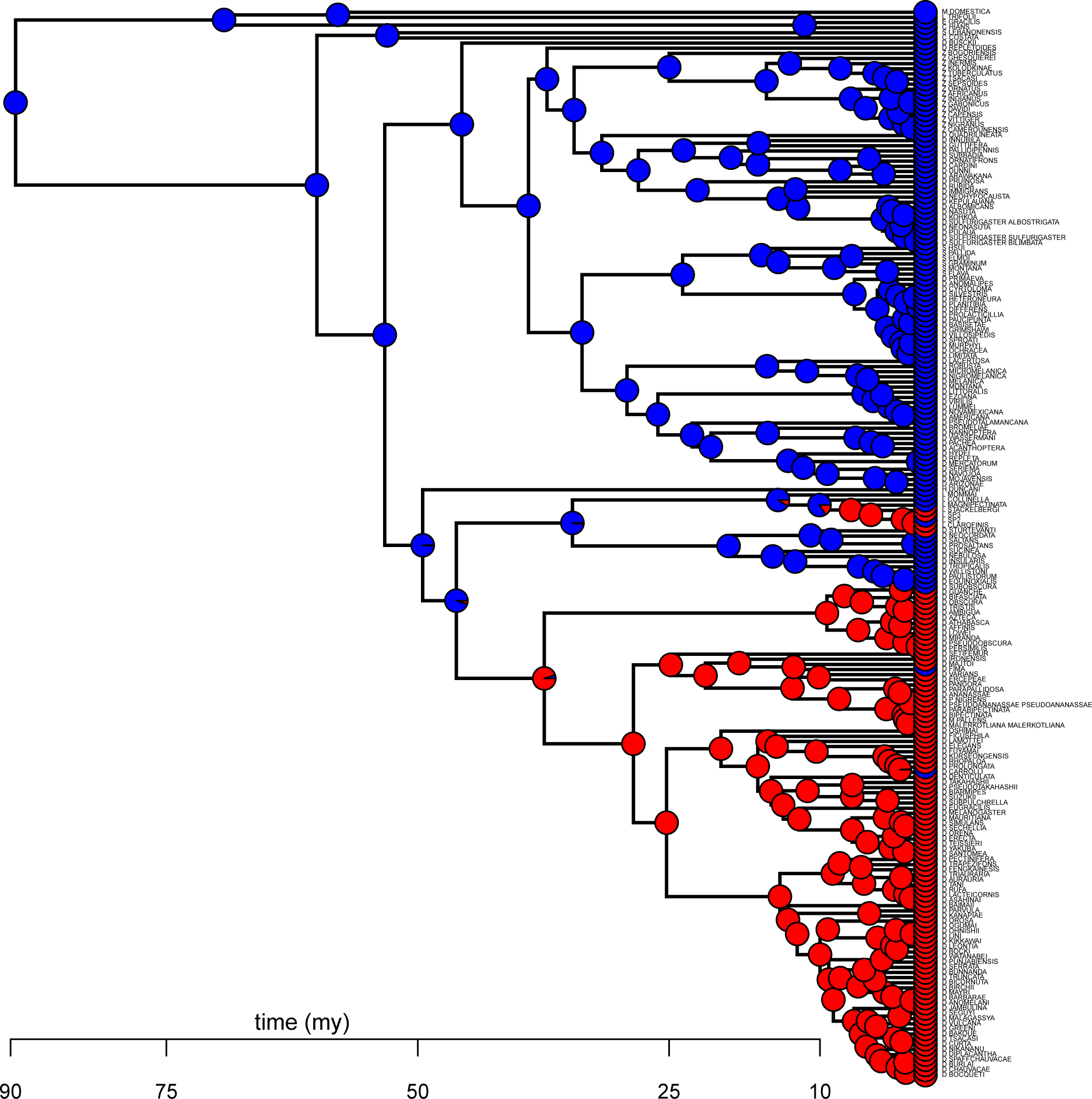
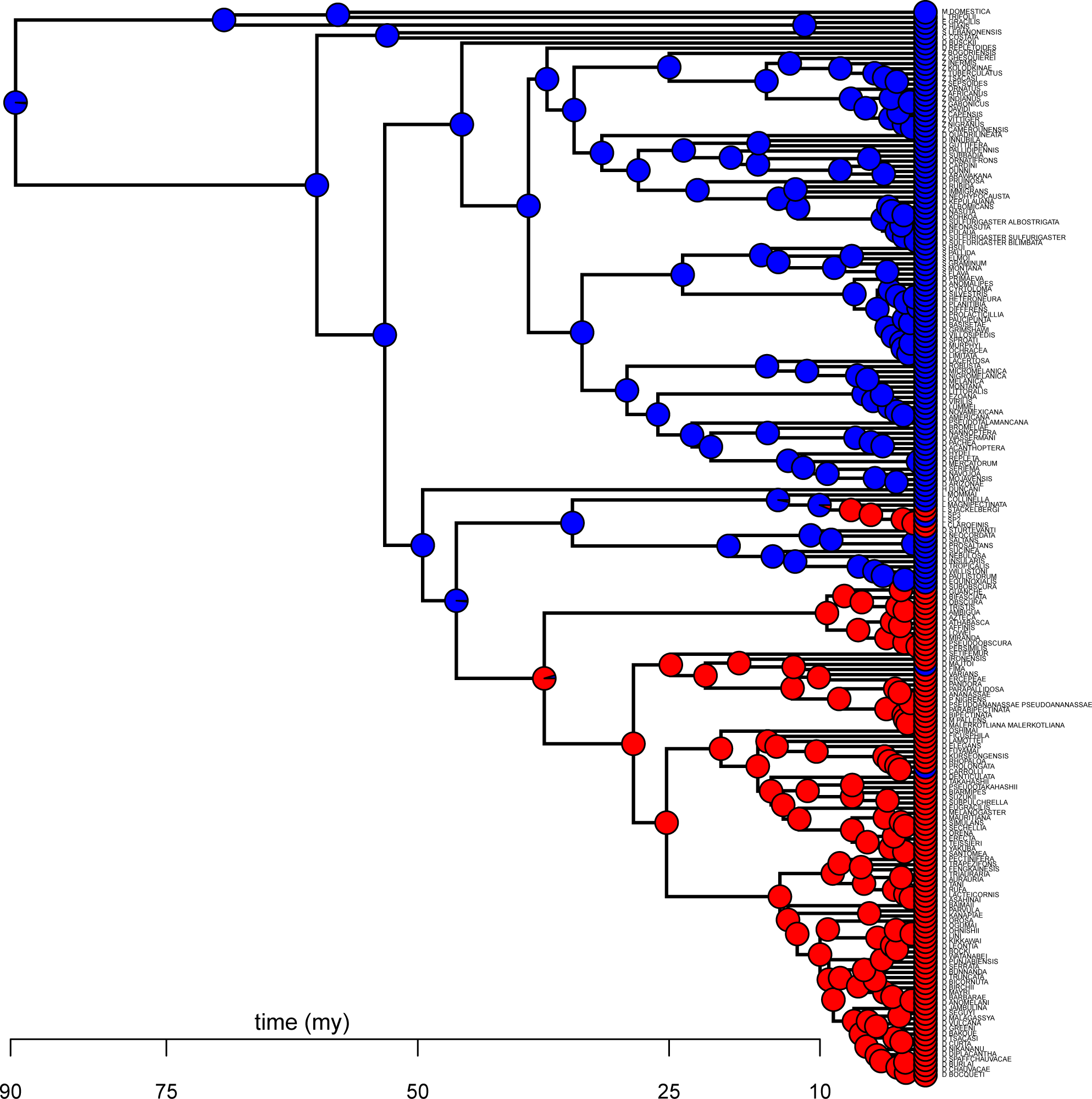
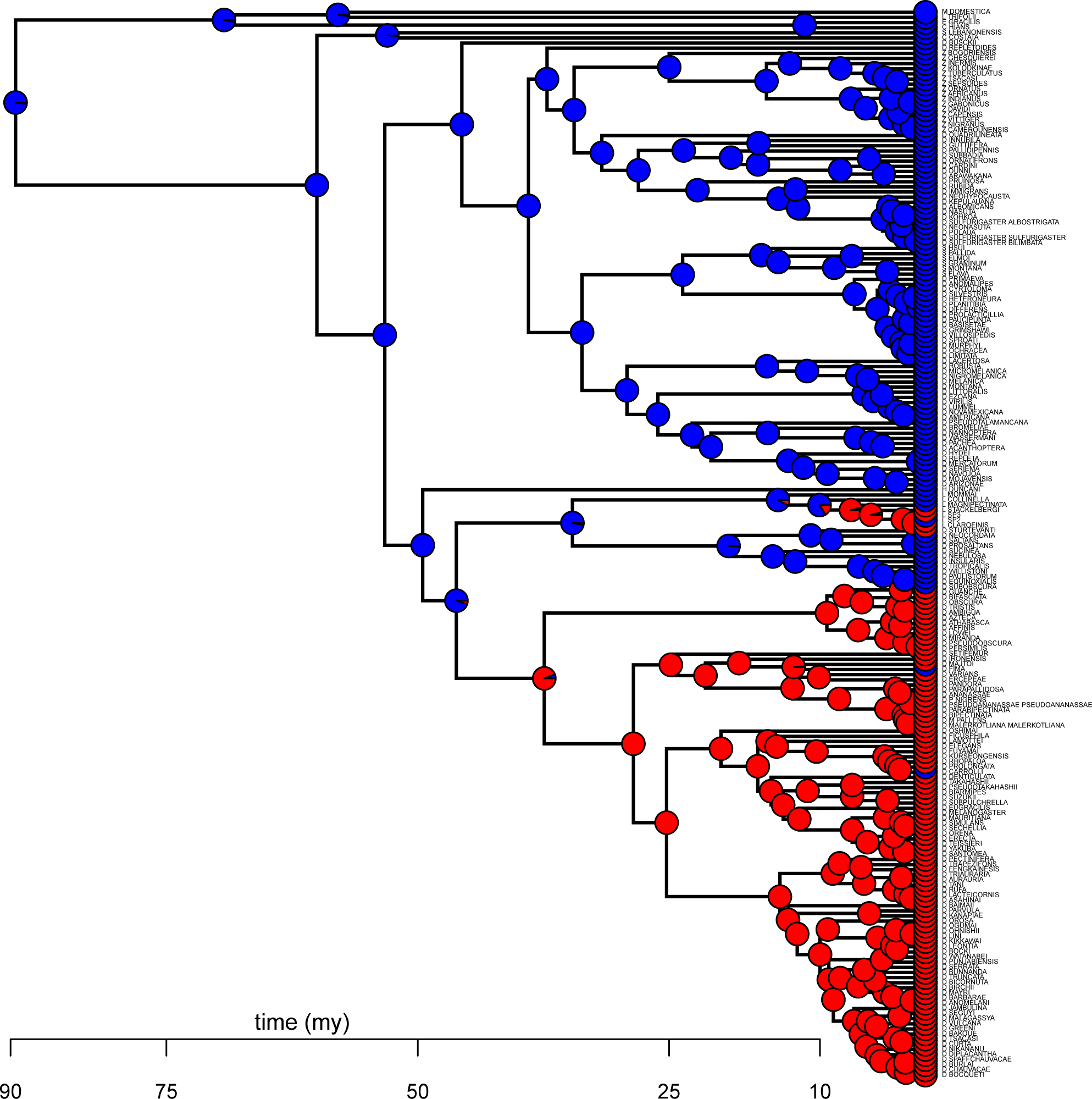
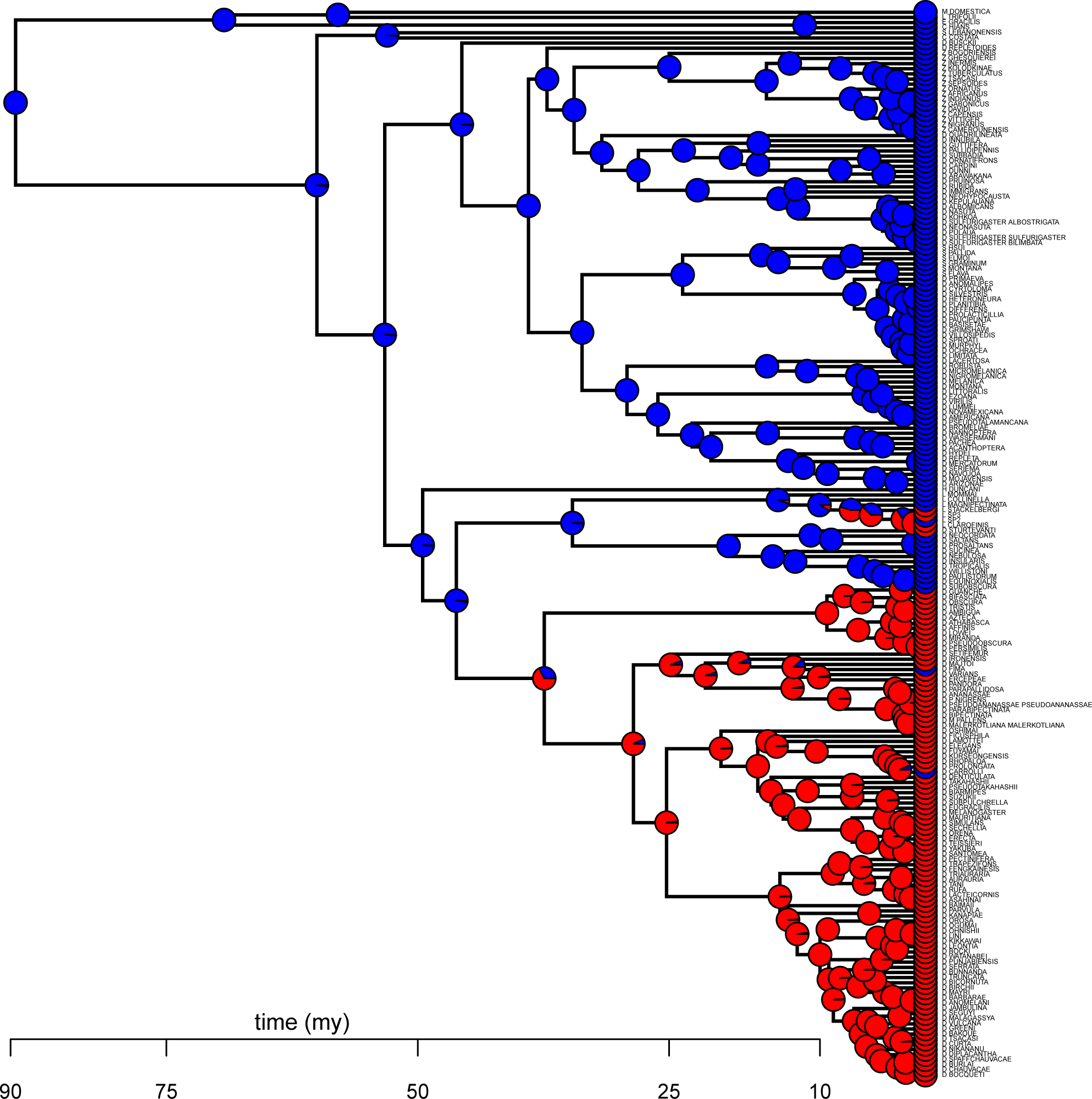
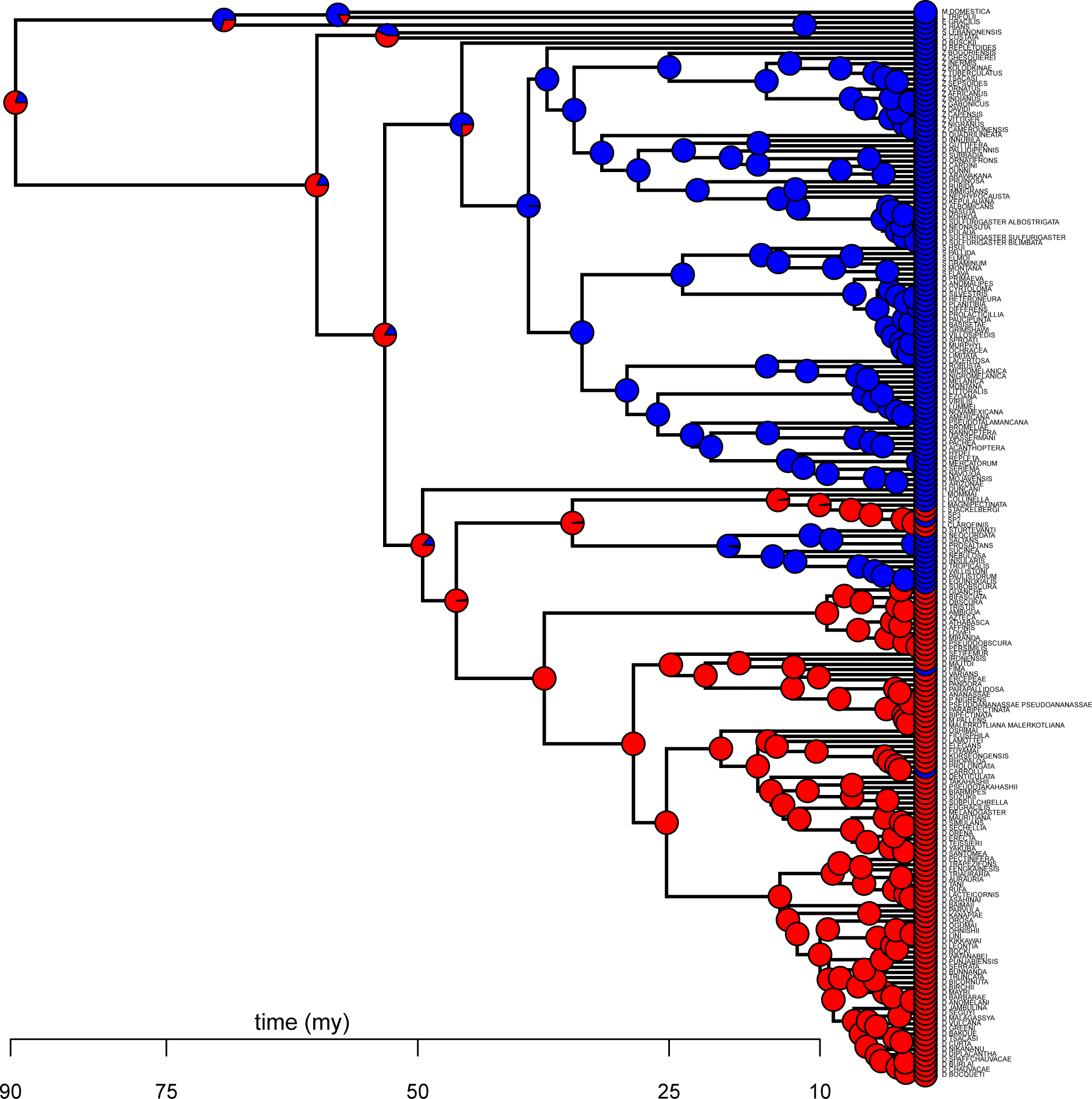
Ancestral state reconstruction under five different models of trait evolution. The phylogenetic tree is based on 250 single-copy BUSCO genes extracted from whole genome sequences (Fig. S4A). Species are colored according to sex comb state (red = present, blue = absent). Pie charts at internal nodes reflect estimated probabilities of ancestral character states under five different models of trait evolution: (A) MK model with unequal rates and a strict molecular clock (Lewis, 2001); (B) MK model with unequal rates and a random local relaxed clock (Drummond and Suchard, 2010); (C) Hidden states variable rates model with two latent rate classes (Beaulieu et al., 2013); (D) Modified threshold model (Felsenstein, 2005) with 9 ordered latent states; (E) Approximation of a Dollo model, with rate of loss >300-fold higher than rate of gain. See Methods and Table S5 for details of ancestral character reconstruction.

**Figure S6.**
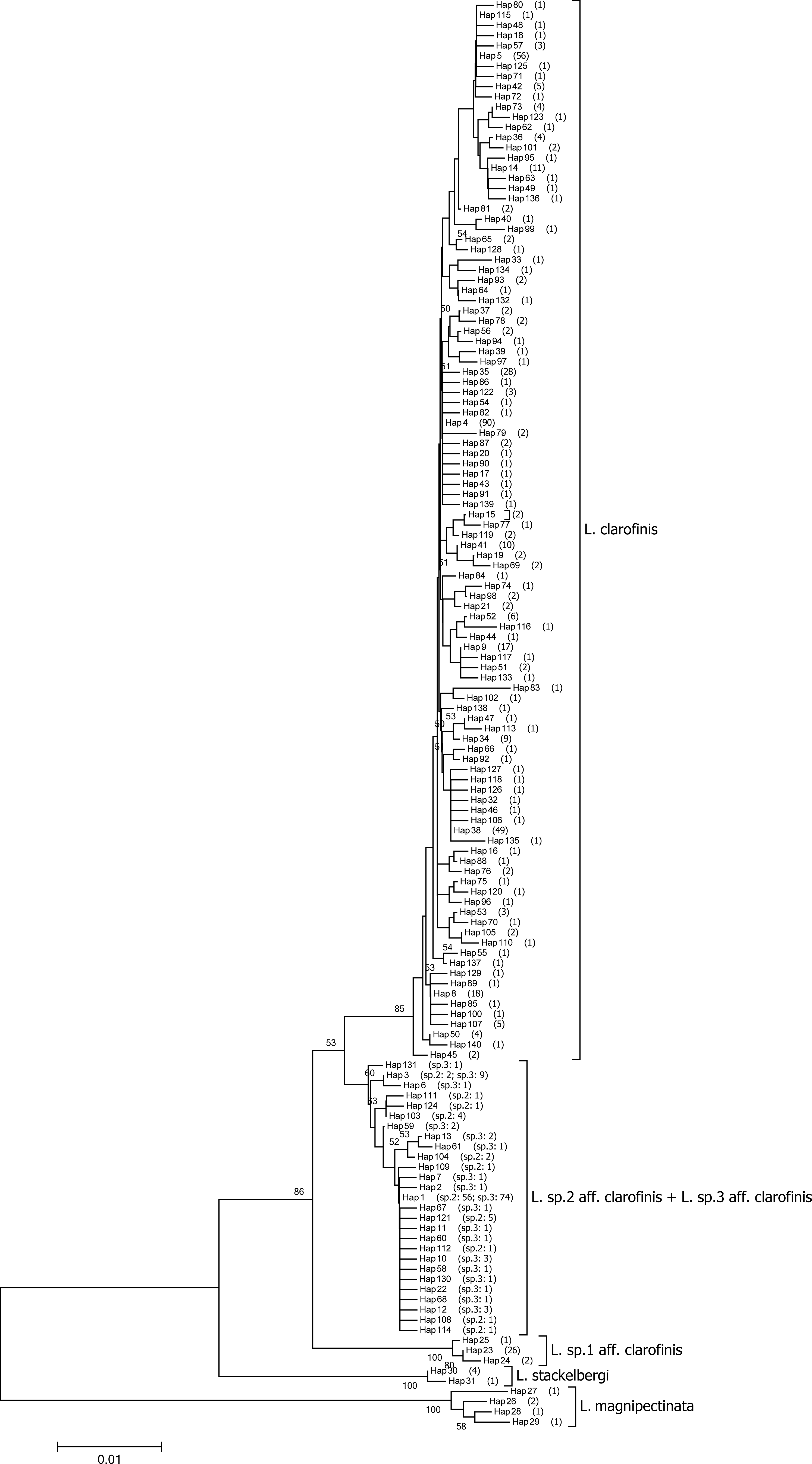
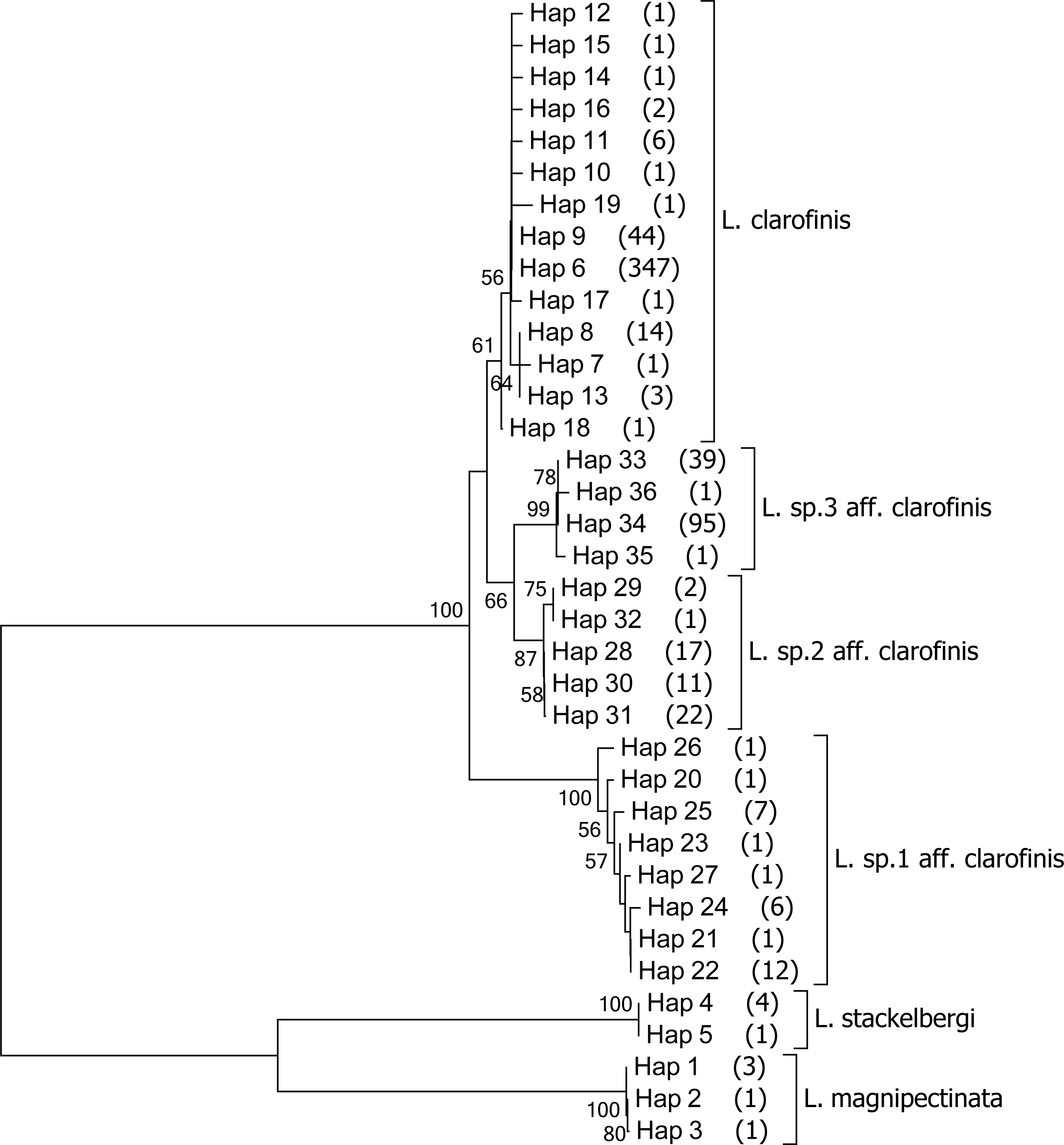
Haplotype trees of the *L. clarofinis* species complex. (A) Unrooted NJ tree of 140 selected *COI* haplotypes. For each haplotype, the frequency is given in parentheses (with the name of the corresponding species indicated in the cluster of *L.* sp.2 aff. *clarofinis* + *L.* sp.3 aff. *clarofinis*). Note that haplotypes 1 and 3 are both shared by these two species. (B) Unrooted NJ tree of all 36 ITS1 haplotypes. Frequency is given in parentheses for each haplotype.

**Figure S7.**
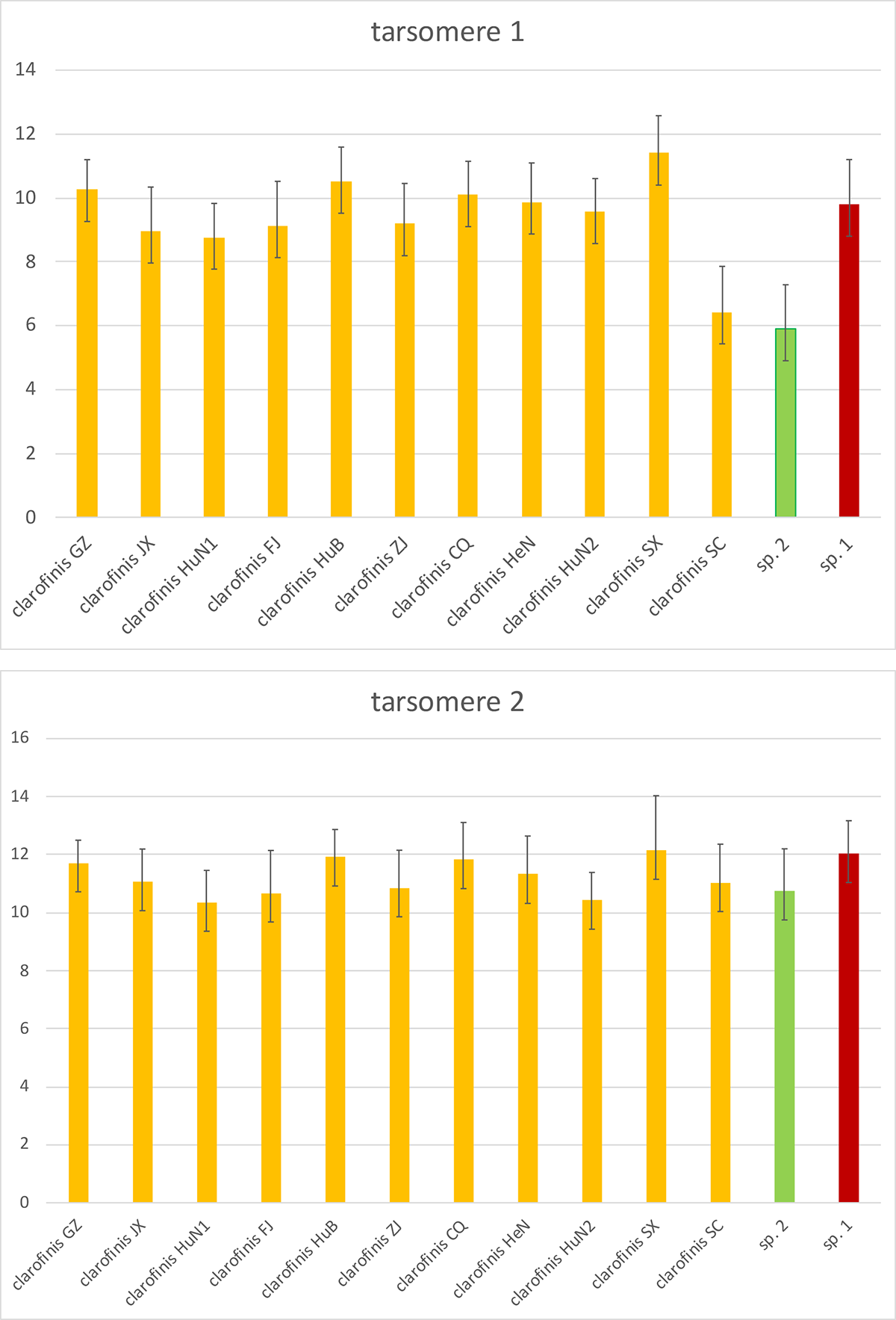
Sex comb size (number of sex comb teeth) in the *clarofinis* complex. Species are coded by color: yellow = *L. clarofinis*, green = sp.2, red = sp.1. All strains of sp.3 lack sex combs. For *L. clarofinis*, different geographic populations are shown separately. Note that *L. clarofinis* population from Sichuan (SC) is more similar to sp.2 than to other *L. clarofinis* populations in the number of sex comb teeth on the first (most proximal) tarsal segment.

