## Supplemental table 5 for "Secondary reversion to sexual monomorphism associated with tissue-specific loss of *doublesex* expression"

Table S5. Prior distribution assumptions for model parameters used for ancestral character reconstruction.

* For all models, transition rate matrix is F81 model which is parameterized by the stationary frequency of the states; q ~ Dirichlet(𝛼_i_,...,𝛼_K_) where K is number of states.

| MK unequal rates | Clock rate  Root frequency | r  𝜋 | ~ Exponential(10)  = [0.5, 0.5] |
| --- | --- | --- | --- |
| RLC | Root frequency  Expected number of rate shifts  Probability of no rate shift on branch i  log Mean branch rate  Standard deviation of branch rates  Rate shift factor on branch i | 𝜋  n  𝜌_i_  log_10_ 𝜇  𝜎  𝜓_i_ | = [0.5, 0.5]  = 2  = n/num_branches = 0.995  ~ Uniform(-5, 2)  ~ Exponential(1/0.587)  ~ 𝜌_i_ + (1 - 𝜌_i_)logNormal(-𝜎^2^/2, 𝜎) |
| Hidden rates class model | Root frequency  Transiton rates  log mean branch rate | 𝜋_i_  q  log_10_ 𝜇 | = 0.25 ∀ i ∈ {1,...,4}  ~ Dirichlet(1,1,1,1)  ~ Uniform(-10,2) |
| Threshold model | Root frequency  Transition rates  log mean branch rate | 𝜋_i_  q  log_10_ 𝜇 | = 0.1 ∀ i ∈ {1,2,...,10}  ~ Dirichlet(⅓, ⅓, …, ⅓)  ~ Uniform(-10,2) |
| Approximate dollo | Root frequency  Transition rates  log mean branch rate | 𝜋  q  log_10_ 𝜇 | = [0.5, 0.5]  ~ Dirichlet(1000, 1)  ~ Uniform(-20, 2) |
